# Spatial drivers and pre-cancer populations collaborate with the microenvironment in untreated and chemo-resistant pancreatic cancer

**DOI:** 10.1101/2021.01.13.426413

**Authors:** Daniel Cui Zhou, Reyka G. Jayasinghe, John M. Herndon, Erik Storrs, Chia-Kuei Mo, Yige Wu, Robert S. Fulton, Matthew A. Wyczalkowski, Catrina C. Fronick, Lucinda A. Fulton, Lisa Thammavong, Kazuhito Sato, Houxiang Zhu, Hua Sun, Liang-Bo Wang, Yize Li, Chong Zuo, Joshua F. McMichael, Sherri R. Davies, Elizabeth L. Appelbaum, Keenan J. Robbins, Sara E. Chasnoff, Xiaolu Yang, Ruiyang Liu, Ashley N. Reeb, Michael C. Wendl, Clara Oh, Mamatha Serasanambati, Preet Lal, Rajees Varghese, R. Jay Mashl, Jennifer Ponce, Nadezhda V. Terekhanova, Nataly Naser Al Deen, Lijun Yao, Fang Wang, Lijun Chen, Michael Schnaubelt, Sidharth V. Puram, Albert H. Kim, Sheng-Kwei Song, Kooresh I. Shoghi, Tao Ju, William G. Hawkins, Ken Chen, Deyali Chatterjee, Hui Zhang, Milan G. Chheda, Samuel Achilefu, David G. DeNardo, Stephen T. Oh, Feng Chen, William E. Gillanders, Ryan C. Fields, Li Ding

## Abstract

Pancreatic Ductal Adenocarcinoma (PDAC) is a lethal disease with limited treatment options and poor survival. We studied 73 samples from 21 patients (7 treatment-naïve and 14 treated with neoadjuvant regimens), analyzing distinct spatial units and performing bulk proteogenomics, single cell sequencing, and cellular imaging. Spatial drivers, including mutant *KRAS*, *SMAD4*, and *GNAQ,* were associated with differential phosphosignaling and metabolic responses compared to wild type. Single cell subtyping discovered 12 of 21 tumors with mixed basal and classical features. Trefoil factor family members were upregulated in classical populations, while the basal populations showed enhanced expression of mesenchymal genes, including *VIM* and *IGTB1*. Acinar-ductal metaplasia (ADM) populations, present in 95% of patients, with 46% reduction of driver mutation fractions compared to tumor populations, exhibited suppressive and oncogenic features linked to morphologic states. We identified coordinated expression of TIGIT in exhausted and regulatory T cells and Nectin receptor expression in tumor cells. Higher expression of angiogenic and stress response genes in dendritic cells compared to tumor cells suggests they have a pro-tumorigenic role in remodeling the microenvironment. Treated samples contain a three-fold enrichment of inflammatory CAFs when compared to untreated samples, while other CAF subtypes remain similar. A subset of tumor and/or ADM-specific biomarkers showed differential expression between treatment groups, and several known drug targets displayed potential cross-cell type reactivities. This resolution that spatially defined single cell omics provides reveals the diversity of tumor and microenvironment populations in PDAC. Such understanding may lead to more optimal treatment regimens for patients with this devastating disease.

**HIGHLIGHTS:**

1. Acinar-ductal metaplasia (ADM) cells represent a genetic and morphologic transition state between acinar and tumor cells.
2. Inflammatory cancer associated fibroblasts (iCAFs) are a major component of the PDAC TME and are significantly higher in treated samples
3. Receptor-ligand analysis reveals tumor cell-TME interactions through NECTIN4-TIGIT
4. Tumor and ADM cell proteogenomics differ between treated and untreated samples, with unique and shared potential drug targets

## INTRODUCTION

Pancreatic ductal adenocarcinoma (PDAC) has a 9% five-year survival rate (Siegel et al., 2020). This poor prognosis is due to early metastases, late detection, and therapy resistance; at diagnosis, the cancer is often unresectable, locally advanced, and/or metastatic disease (McGuigan et al., 2018). The incidence of pancreatic cancer has increased in recent years, with a roughly 5% rate increase in the last decade, which is projected to raise PDAC from the 4th to the 2nd most common cause of cancer-related death in the U.S. by 2020 (Saad et al., 2018; Rawla et al., 2019; Ilic & Ilic, 2016; American Cancer Society, 2020). The first-line treatment for pancreatic cancer is surgery, if possible, followed by radiation and/or chemotherapy (Conroy et al., 2018; Kang et al., 2018). Despite the promising successes of immunotherapy in several cancer types, there have been very limited responses to immunotherapy in PDAC (Morrison et al., 2018; Balachandran et al., 2019). Nearly all patients will develop chemoresistant tumors and develop progressive, metastatic PDAC within two years of diagnosis, and beyond the two FDA-approved chemotherapy regimens (FOLFIRINOX and gemcitabine+nab-paclitaxel), there are no effective treatment regimens.

Over the last decade, there have been several major efforts to characterize the genomic and transcriptomic landscape of PDAC (Raphael et al., 2017; Moffitt et al., 2015; Collisson et al., 2011; Bailey et al., 2016). Altered *KRAS, TP53, CDKN2A,* and *SMAD4*, among others, have been identified as key disease drivers, with KRAS hotspot mutation rates as high as 97% in some cohorts (Raphael et al., 2017). Several expression-based subtyping strategies have been developed. Moffitt and colleagues classified tumors into classical or basal-like subtypes based on their expression profile; this is the most widely applied system (Moffitt et al., 2015). However, a major challenge in bulk sequencing analyses of PDAC is that tumor samples contain low neoplastic cellularity due to the presence of high amounts of a dense desmoplastic stroma composed of cells that affect the ability to target and treat PDAC. While several approaches to address low tumor purity have been applied, including ultra-high depth sequencing and microdissection strategies, low neoplastic cellularity remains a significant challenge for data analysis and interpretation of sequencing studies in PDAC (Raphael et al., 2017; Moffitt et al., 2015; Maurer et al., 2019).

Single cell technologies are well equipped to address the low tumor cellularity challenge in PDAC by enabling tumor cell analysis regardless of tumor content of a given sample. Additionally, single cell level resolution allows for the dissection and analysis of different cell types in the tumor microenvironment (TME) and their interactions with tumor cells. This addresses another challenge in the field. We know that non-tumor cell components of the TME play a critical role in PDAC, yet the mechanistic underpinnings behind the TME’s role in PDAC is largely unknown. For instance, several cancer associated fibroblast (CAF) subtypes, including myCAFs, iCAFs, and apCAFs, have been identified. Cytotoxic NK and CD8+ T cells are downregulated (Elyada et al., 2019; Sahai et al., 2020; Schnurr et al., 2015; Looi et al., 2019). This creates an immunosuppressed, pro-tumorigenic environment, but how this occurs is poorly understood (Uzunparmak et al., 2019; Ren et al., 2018). Furthermore, tumor populations are seldom uniform and single cell technologies allow for an in-depth analysis of cellular heterogeneity (Alizadeh et al., 2015; Chung et al., 2017).

Herein, we use a spatially distinct, multi-sampling approach to analyze 73 PDAC samples across 21 patients who have undergone different treatment regimens (treatment-naïve, neoadjuvant FOLFIRINOX, neoadjuvant gemcitabine+nab-paclitaxel, and Chemo-RT). This spatial approach allowed us to interrogate both inter- and intra-tumor heterogeneity via extensive omics, including bulk DNA and RNA sequencing, bulk proteomics and phosphoproteomics, single cell RNA sequencing (scRNA-Seq), and cellular imaging.

## RESULTS

### Study Design and Single Cell Overview of the Cohort

We collected 73 pancreatic ductal adenocarcinoma samples from 21 patients undergoing standard treatment, including 4 normal adjacent tissue (NAT) samples. The various treatment groups included 7 treatment-naïve cases, 8 neoadjuvant FOLFIRINOX cases, 4 neoadjuvant gemcitabine+nab-paclitaxel cases, 1 mixed (FOLFIRINOX and gemcitabine+nab-paclitaxel), and 1 Chemo-RT case. Each tumor was spatially sampled 2–4 times, with sample segments subsequently used to generate histologic, imaging, and omics data, H&E slides, imaging mass cytometry (IMC), single cell RNA-Sequencing (scRNA), bulk mass spectrometry-based proteomics and phosphoproteomics, bulk whole exome sequencing (WES), and bulk RNA sequencing (RNA-Seq) (**Figure 1A, Table S1, STAR Methods**). We generated scRNA data for all 73 samples, WES for 64 samples, and bulk RNA-Seq for 65 samples. A subset of our cohort (n = 30) underwent TMT11 proteomic and phosphoproteomic characterization. We generated IMC slides using the Hyperion imaging system for 12 samples from 4 cases, each with 3–4 regions of interest (ROI) (**Figure 1B**).

**Figure 1:**
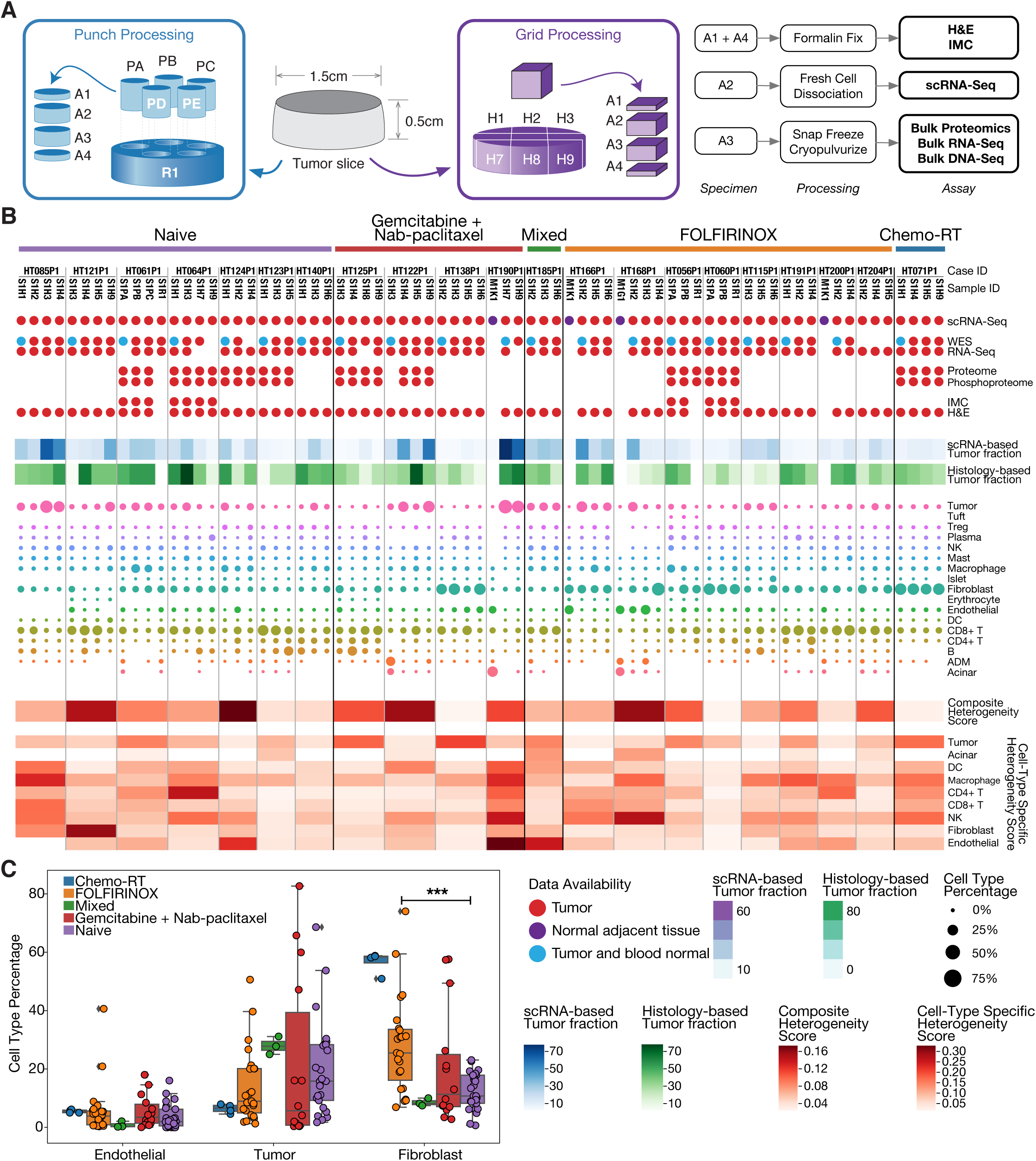
Sampling Strategy and Cohort Overview. A) Spatial sampling approach. At least 2 punches or grids were selected from each tumor for comprehensive imaging and omics characterization. “P” denotes tissue punches, “H” denotes tissue grids, and “R” denotes remainder tissue. **B)** Top: Data overview of the cohort. M1K1 and M1G1 denote NAT samples. Middle: scRNA-based and histology-based estimates of tumor purity and scRNA-based cell type percent composition. The size of each circle denotes percentage. Bottom: Composite cell type proportion and expression heterogeneity score (**STAR Methods**). A higher score denotes higher heterogeneity. Cell type-specific scores are calculated in a similar fashion but are based only on expression. **C)** Endothelial cell, tumor cell, and fibroblast percentages in samples split by treatment group.

Following QC, we scaled and normalized scRNA data, clustered more than 230k cells across all samples based on expression profiles, and assigned cell types based on marker gene expression (**Figure S1A, STAR Methods**). Cell types included: acinar, acinar ductal metaplasia (ADM), B, CD4+ and CD8+ T cells, dendritic cells (DCs), endothelial cells, erythrocytes, fibroblasts, islet cells, macrophages, mast, natural killer (NK), plasma cells, naïve and regulatory T cells, tuft cells, and tumor cells (**Figures 1B and S1B**). Using the fraction of tumor cells as a proxy for tumor purity, we estimated that tumor purities ranged from 0.12% to 82.68% across samples, with an average of 17.16%.

We calculated a composite heterogeneity score to quantitatively determine cell-type proportion and expression differences across cells between the spatial samples within each tumor (**Figure 1B, STAR Methods**). Briefly, this score is based on mean pairwise correlation of global average scRNA expression and the cell type proportions between samples from the same case. Samples from treated patients tended to have significantly higher percentages of stroma cells (p = 0.001) (**Figures 1B and S1C**). In particular, FOLFIRINOX samples had significantly higher percentages of fibroblasts (p = 10^−4^) (**Figure 1C**). We did not observe significant endothelial or tumor cell number differences between treatment groups. ADM cells were present in 20/21 cases but were most abundant in HT122P1 (treated with gemcitabine+Nab-paclitaxel) and HT168P1 (FOLFIRINOX). More than half the cases had high degrees of spatial heterogeneity (composite score > 0.1) regardless of treatment status, suggesting that cell type proportions and tumor subpopulations have substantial spatial variation. Pathologic review of H&E slides revealed that within-patient tumor content differences across samples averaged 24%, with a range of 5% to 64% (**STAR Methods**), consistent with tumor percentages from scRNA (Pearson R=0.36, p=0.003). To quantify spatial heterogeneity at a cell-type level, we used a similar expression-based approach for each individual cell type (**Figure 1B, STAR Methods**). The greatest heterogeneity occurred amongst tumor cells, followed by macrophages and endothelial cells, suggesting our spatial sampling captures differences in tumor subpopulations within each case, as well as those within stromal and immune fractions. Principal component analysis (PCA) of bulk proteomic and phosphoproteomic data confirm that, while most within-tumor regions cluster close to one other, several specimens from the same tumor are quite different, representing intra-tumor heterogeneity (**Figures S1D and S1E**). These results underscore limitations of a strict bulk-data approach, in which such heterogeneity may remain invisible.

### Genomic Landscape and Single Cell Subtyping Heterogeneity

Using paired bulk whole exome sequencing (WES), we investigated somatic and germline variants, observing substantial variation of driver mutation variant allele fractions (VAFs) between samples (**Figure 2A, Table S2**). We detected a KRAS hotspot variant in at least one sample in all cases (HT204P1 has no WES data). Furthermore when considering mutations with extremely low VAFs ( < 0.01), 10 of the 20 cases had samples whose mutation profiles differed from one another. While it is possible these mutations existed in all samples in a given case and that detection was imperfect, materials were sequenced and processed uniformly and data underwent extensive QC and manual genotyping, suggesting this difference was not due to technical shortcomings (**STAR Methods**). We also characterized germline variants using CharGer in order to identify pathogenic and likely pathogenic variants in the cohort (Scott et al., 2019) (**STAR methods**). Three cases carried such variants in the homology-directed DNA repair pathway (*FANCC* p.D23\**, BRCA2* p.I1470* and p.K607*, and *ATM* p.Y1124*), and as expected, all samples carried the same variant in each case.

**Figure 2:**
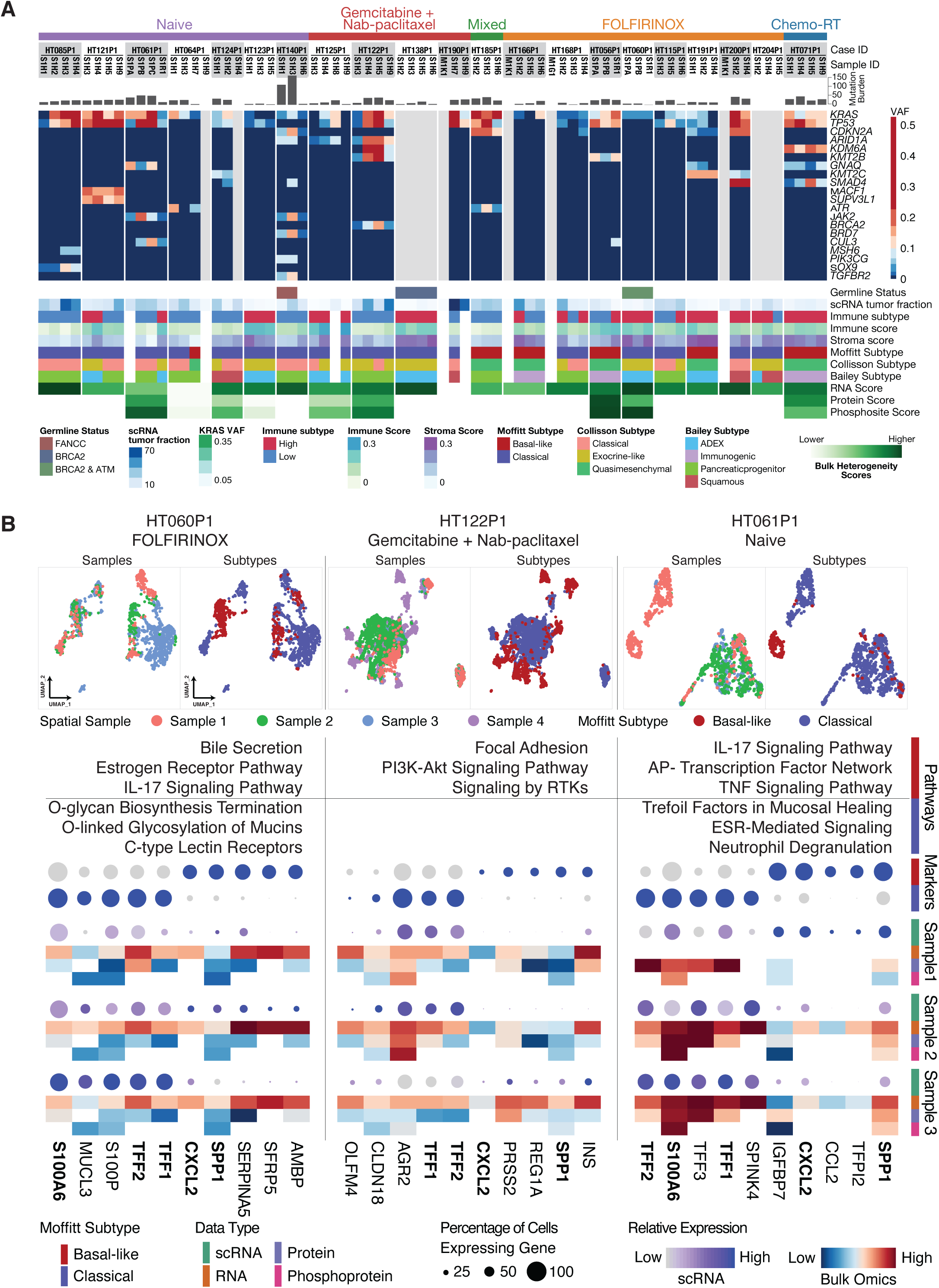
Genomic Landscape and Single-Cell Subtyping. A) Top: Genomic landscape of the cohort showing the top significantly mutated genes. Color scale denotes VAF for each gene. Bottom: Bulk omics overview of the cohort. **B)** Three examples of cases with heterogeneous tumor cells with different single-cell subtypes. UMAP plots show only clustered tumor cells. For each case, the top five significant genes ranked by fold change across subtypes are shown. Bolded genes represent genes that are present across at least two of the cases.

Using bulk RNA-Seq, we classified each sample into previously reported subtypes, including those defined by Moffitt et al. (2015), Collisson et al. (2011), and Bailey et al. (2016) (**Figure 2A**) and determined immune subtypes and stromal and immune compartment scores using xCell and ESTIMATE, respectively (Yoshihara et al., 2013; Aran et al., 2017) (**STAR Methods**). Aside from 1 case, Moffitt subtyping was consistent within-tumor across samples, while Collisson and Bailey subtyping both showed larger numbers of subtype heterogeneity across samples. We observed differences in immune subtypes between spatial samples in 7 tumor cases (**Figure 2A**). We also observed a weak positive correlation between scRNA stroma percentages and ESTIMATE stroma scores (Spearman R: 0.46, p = 10^−4^) though not between scRNA immune fraction percentages and ESTIMATE immune scores (**Figure S2A and S2B**).

In order to further dissect patient-specific heterogeneity, we selected three cases representing our major treatment groups, HT060P1 - FOLFIRINOX, HT122P1 - gemcitabine + nab-paclitaxel, and HT061P1 - treatment-naïve, for further analysis. Given the consistency of Moffitt subtyping, we used classical and basal-like expression profiles to classify tumor cells at single cell resolution (Torres & Grippo et al., 2018) (**Figures 2B and S2C, STAR Methods**). Specifically, we evaluated tumor-only clusters and annotated them by their spatial sample origin and their single cell Moffitt subtype for each case. In all three cases, there are both mixed subpopulations (originating from > 1 sample) or sample-specific subpopulations, representing spatially distinct tumor clusters. Interestingly, we also observe discrete clusters separated by subtype and these tend likewise to be of different spatial origin. In HT061P1, the basal-like cluster does not correspond to any mappable *KRAS* mutations (**Figure S2C and S2D**). Differential gene expression (DGE) analysis shows that basal-like tumor cells are enriched in IL-17 and PI3K-AKT signaling and AP-transcription factor network pathways. Classical tumor cells are enriched in O-linked glycosylation of mucins and trefoil factors in mucosal healing pathways (**Figure 2B**). This suggests that spatial sampling captures spatially distinct tumor cells having different expression profiles that likely correspond to different subtypes within the same patient. Extending this analysis to the rest of the tumor cohort, we identified substantial amounts of mixed classical and basal-like tumor cells in 12/21 patients (defined as > 10% of tumor cells in the less common subtype) and determined that the basal tumor cells upregulate a number of EMT genes, including VIM and *ITGB1* (**Figure S2E**).

Using scRNA analysis, we identified the top 5 most differentially expressed genes (DEGs) between classical and basal-like cells from each sample within a tumor and followed with the complementary analysis using bulk data. Across the above three within-patient tumor specimens, we identified consistent scRNA DEGs between classical and basal tumor cells, including *S100A6*, *TFF1*, and *TFF2* in classical cells and *SPP1* and *CXCL2* in basal-like cells (**Figure 2B**). Spatially distinct samples enriched in classical or basal-like tumor cells strongly express these subtype-specific DEGs and their expression patterns are consistent across omics data types (**Figure 2B**). For instance, in case HT122P1, sample 1 (S1H3) and sample 2 (S1H4) express classical markers at the scRNA, bulk RNA, protein, and phosphoprotein levels. Sample 3 (S1H5) expresses both classical and basal-like markers at the scRNA and RNA levels, with slightly higher levels of basal-like expression. In short, this analysis demonstrates it is clearly possible to identify distinct tumor subpopulations in space within a tumor. The data here indicate PDAC tumors are heterogeneous and are not readily classifiable as a single subtype and this may explain, in part, the intrinsic and acquired resistance observed in PDAC. Therefore, therapeutic decisions that assume homogeneity may miss these underlying nuances.

### KRAS signaling and Spatial Drivers

We reclustered tumor cells from all samples based on scRNA expression to assess their heterogeneity. The majority grouped into patient-specific clusters, consistent with the varying genetic backgrounds and alterations in each patient (**Figure S3A**). We also mapped mutations called from the bulk whole exome data to single cells (**Figure 3A, STAR Methods**), observing three groups of tumor cells originating from a large number of patients. Two groups had multiple different *KRAS* mutations (**Figures 3A and S3B**). The remaining group is significantly depleted in *KRAS* mutations and likewise lacks *TP53* or *CDKN2A* mutations (**Figure 3B**). We denote these three populations as “mixed” and hypothesize that their consistent expression profiles across samples may signify a potential common element in PDAC.

**Figure 3:**
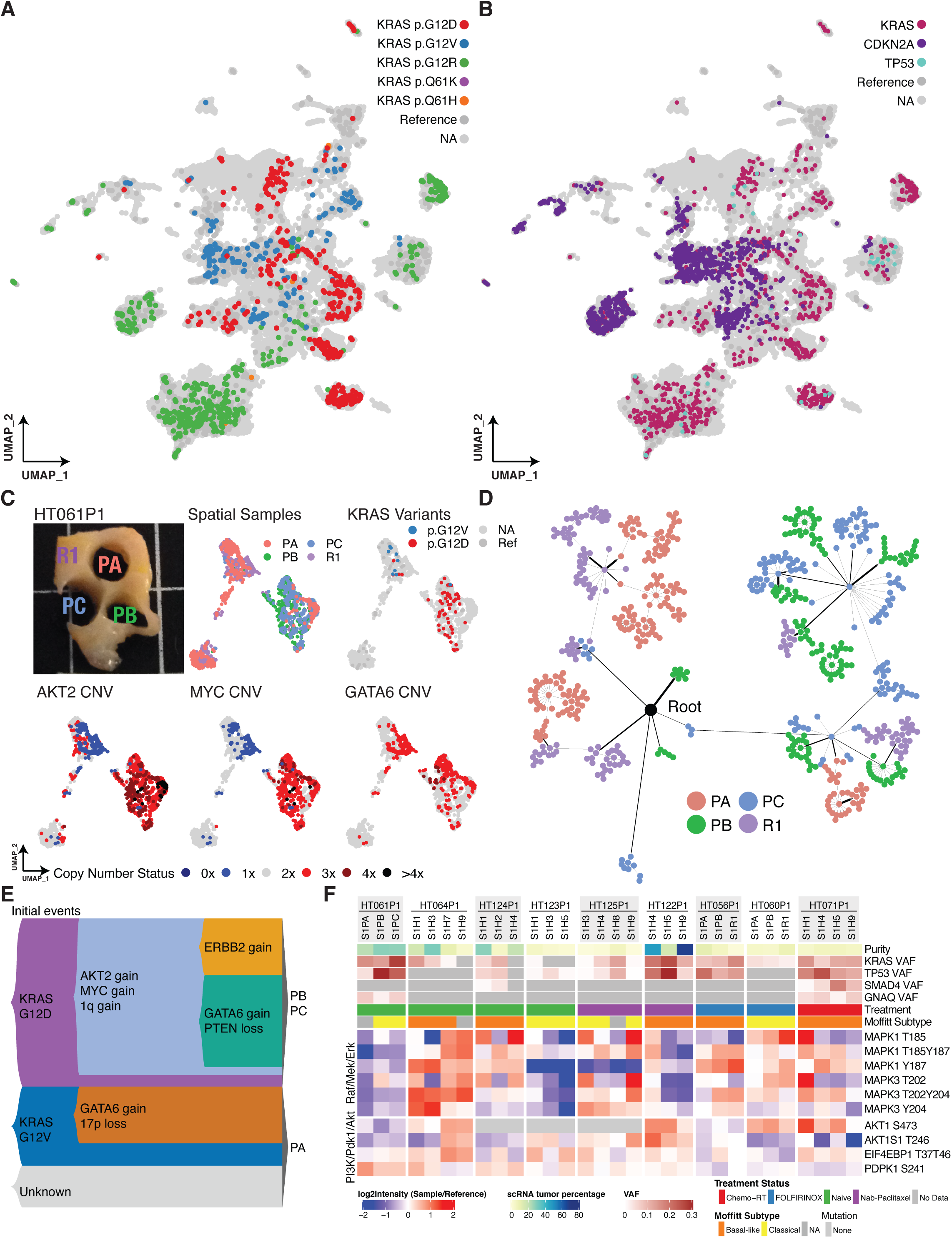
Spatial and Oncogenic Driver Heterogeneity. A) Tumor cell clusters labeled with *KRAS* hotspot mutations. **B)** Tumor cell clusters labeled with *KRAS, CDKN2A,* and *TP53* mutations. **C)** Case HT061P1.. Top row left to right: tissue sample spatial locations, sample IDs (R1 denotes the remainder tissue), *KRAS* variants. Second row left to right: *AKT2* CNV, *MYC* CNV, and *GATA6* CNV. Copy number calls were obtained using inferCNV. **D)** CNV-based lineage tree of a subset of tumor cells from HT061P1. **E)** Proposed model of tumor progression for HT061P1. **F)** Bulk phosphosite levels in the PI3K/Pdk1/Akt and Raf/Mek/Erk pathways. Grey boxes denote missing data. Samples with proteomics/phosphoproteomics did not all have mutations in *CDKN2A*.

We conducted DEG analysis for each of these three mixed tumor populations, finding that *SPP1* and *CXCL2*, two of our previously identified single-cell classical subtype markers, were upregulated in the *KRAS* WT populations (**Figures 2B and S3C**). Conversely, these genes are expressed at a low level in both the *KRAS* mutated populations. Additionally, in order to test the impact of different *KRAS* hotspot variants, we compared the gene expression profiles of the subset of tumor cells with *KRAS* mutations against each other (**STAR Methods**). Interestingly, we found that compared to other KRAS mutations, tumor cells that harbor *KRAS* p.G12V upregulate several genes associated with more aggressive or metastatic tumors, including *COL1A1*, *VIM*, and *MUC5B* (**Figure S3D**) (Valque et al., 2012; Niknami et al., 2017; Zhang et al., 2018).

As most pancreatic cancers carry a hotspot *KRAS* driver mutation, we additionally manually genotyped common KRAS mutations at the p.G12 and p.Q61 loci (**STAR Methods**). Intriguingly, we identified 5 cases with multiple *KRAS* hotspot drivers, which we denote as cases with multiple *KRAS* clones (**Figure S3G**). Focusing on case HT061P1, we obtained 4 subpopulations when clustering tumor cells, three small clusters largely derived from punch A and one large cluster that was common to all three punches (the remainder, as expected, represented all clusters) (**Figures 3C and S3H**). Notably, we almost perfectly map *KRAS* p.G12V cells into one cluster from punch A predominant populations and p.G12D cells onto the large mixed cluster. Not only do we observe two discrete clones carrying different *KRAS* driver mutations in the same patient, but we also find they were spatially separated and have distinct gene expression profiles (**Figure 3C**). It is uncertain whether the other two clusters from which we did not map a *KRAS* mutation are truly *KRAS* WT due to a paucity of cells.

Using inferCNV, we determined the copy number profile of the tumor cells and identified several different CNV signatures at both focal and arm levels unique to the two differing *KRAS* subclones in case HT061P1 (**Figures 3C and S3I, STAR Methods**) (Tickle et al., 2019). For instance, the p.G12D population has deep amplifications of *AKT2* and *MYC* while both p.G12D and p.G12V clusters harbor amplifications in *GATA6*, among others (**Figures 3C and S3I**). Furthermore, we used the inferred single cell copy number profiles in order to reconstruct a lineage tree using the MEDALT algorithm (**Figure 3D**) (Wang et al., 2020). Interestingly, the CNV-based tree separates the tumor cells into two major groups, consistent with the gene expression-based clustering as well as the spatial origin of the cells (**Figure 3E**). Consolidating this information, we propose a model that integrates the gene expression and CNV data (**Figure 3E**). We observe two tumor subpopulations in punch A, one of which has an unknown initial driver (lack of mappable *KRAS*) and the other is driven by *KRAS* p.G12V. The *KRAS* p.G12V population then acquired an amplification in *GATA6* and a 17p deletion. In punches B and C (and a portion of A), the initial driver was *KRAS* p.G12D. This was followed by a gain of *AKT2*, *MYC*, and 1q and additional subsets of cells acquired either an *ERBB2* amplification or a *GATA6* amplification and *PTEN* deletion (**Figures 3C, 3E, and S3I**). These results provide an example of the vast degree of tumor heterogeneity in PDAC present at the expression, mutational, and CNV levels with corresponding differences in genomic alterations that may impact tumor growth, progression, and response to treatment.

We determined the impact of mutations on downstream targets by analyzing changes in protein and phosphorylation in several oncogenic pathways (**STAR Methods**). At the protein and phosphoprotein level, we observed that samples carrying a *TP53* mutation had several proteins and phosphosites upregulated in the cell cycle and mismatch repair pathways, including MCM7 and CDK1 (**Figures S3E and S3F**). Interestingly, we also identified observed MKI67, a cell proliferation marker, is upregulated at both the protein and phosphoprotein levels in *TP53* mutants. As expected, several members of the RTK Ras pathway had higher phosphorylation abundance in *KRAS* mutants (**Figure S3F**). As the *KRAS* signaling pathway is uniformly upregulated in almost all pancreatic cancers, we analyzed the abundance of key phosphosites in the KRAS pathway within the context of different *KRAS* mutations and treatment groups (**STAR Methods**). Strikingly, we observe a large degree of differential regulation, both between and within tumors, in several phosphosites within thePI3K/PDk1/Akt and Raf/Mek/Erk pathways (**Figure 3F**). There is an association between p53 mutation status and lower phosphorylation levels in MAPK1, MAPK3, and AKT1, among others, seem to potentially be related to. While samples generally clustered together within the same patient, some cases such as HT125P1 and HT122P1 do not. In HT125P1, two samples do not have a detectable KRAS mutation (S1H3 and S1H9) while the other two have a G12V mutation (S1H4 and S1H8), and these samples seem to segregate accordingly into the higher and lower phosphosignaling groups, suggestive of differential RAS activation even within the same patient. In particular, the MAPK1 T185 phosphosite is differentially regulated between these samples, while MAPK1 Y187, for instance, is uniformly expressed throughout HT125P1. While these patterns did not seem to have a connection to specific KRAS hotspot amino acid changes or treatment status, these results underscore the degree of KRAS signaling heterogeneity in PDAC.

### Acinar-Ductal Metaplasia Populations Transition Between Tumor and Acinar Cells

A prevailing model is that pancreatic cancer arises from acinar cells that undergo acinar to ductal metaplasia (ADM) (Makohon-Moore et al., 2018; Murphy et al., 2013; Kopp et al., 2012). However, this is based on mouse models and the actual role this cell state plays in the development of PDAC remains unknown (Storz, 2017). A major hurdle has been the small numbers of acinar and ADM cells sampled from patients at single cell resolution (Peng et al., 2019; Qadir et al., 2020). We detected ADM populations in 20/21 cases, but focused our analyses to the two cases with the most substantial proportion of ADM cells in our cohort: HT122P1 and HT168P1 (**Figure 4A)**. By mapping mutations, we found that most mutations resided in tumor cells and to a lesser extent, ADM cells (**Figures 4B and 4C**). Consistent with previous studies, we observe ADM initiating mutations in KRAS (from HT122P1) and CDKN2A (from HT168P1) present in ADM populations, which we denote as “ADM Mutated,” representing a more advanced state of ADM (Storz, 2017). While tumor and acinar cells express epithelial and acinar cell markers in a mutually exclusive pattern, both ADM populations express a combination of both, consistent with current understanding that they exist in an intermediate, reversible, state (**Figure 4D**). In order to better describe this expression gradient, we created composite tumor and acinar scores using common tumor (n = 23) and acinar (n = 19) marker genes, respectively (**Figure 4E, STAR Methods**). Mapping these scores onto single cells revealed that the ADM populations harbor heterogeneous mixtures of both ductal and acinar composite signatures (**Figure 4E**).

**Figure 4:**
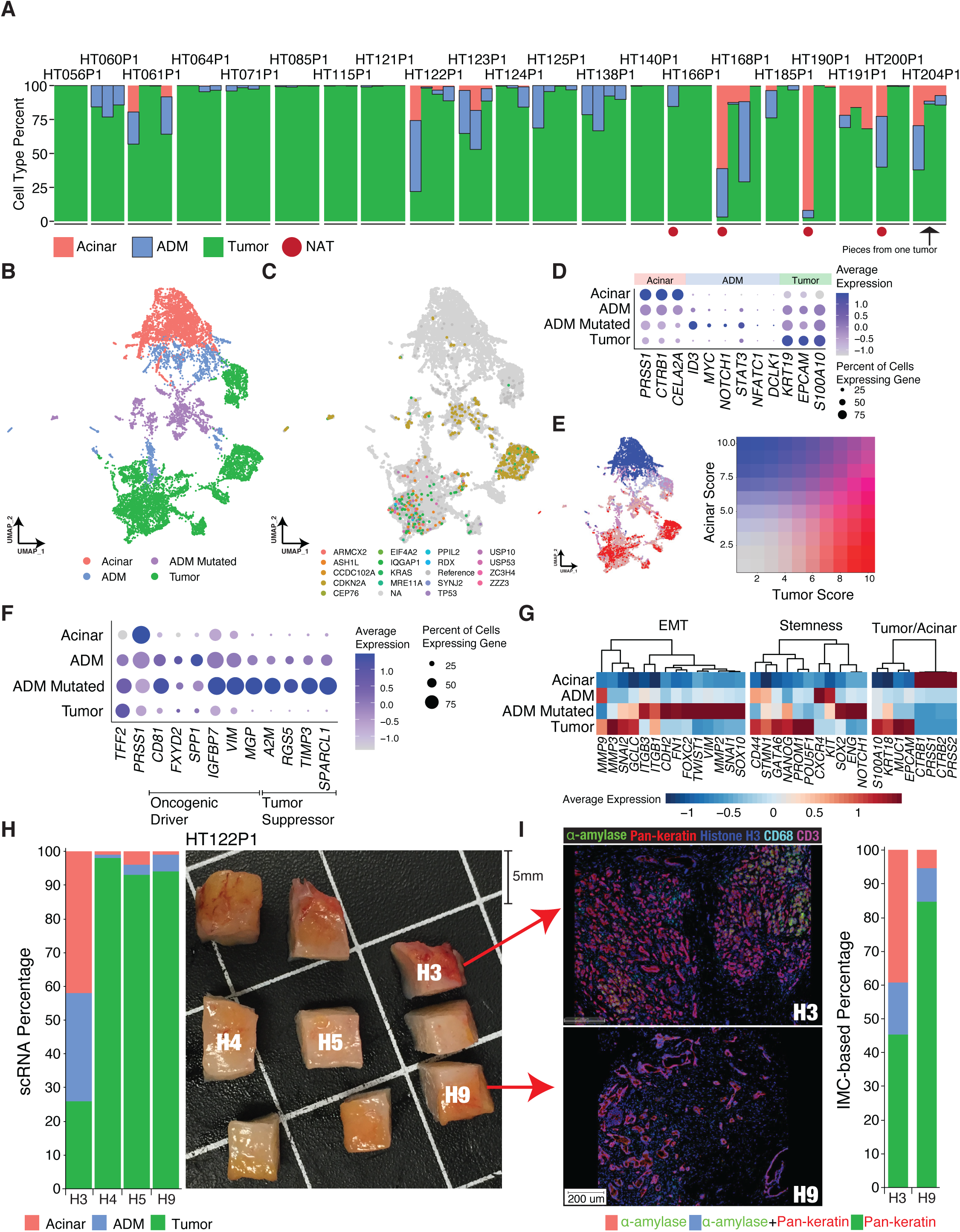
Acinar-Ductal Metaplasia Transition Populations. A) Acinar, ADM, and tumor percentages in all 73 samples. NAT samples are labeled with a red circle. **B)** Acinar, ADM, ADM mutated, and tumor populations from cases HT122P1 and HT168P1. **C)** All mappable mutations in acinar, ADM, and tumor cells of HT122P1 and HT168P1. **D)** A subset of markers used to distinguish between acinar, ADM, and tumor cells. **E)** Composite score using all acinar and tumor markers. **F)** Top significant DEGs in ADM cells. *TFF2* and *PRSS1* are included as references, with each being a strong marker for tumor and acinar cells, respectively. **G)** Average expression of EMT, stemness, and tumor/acinar genes. **H)** scRNA-based acinar, ADM, and tumor cell proportions (*left*) and sample spatial locations (*right*) in samples from HT122P1. **I)** Left: IMC slides (Hyperion) of H3 and H9. The antibody labels used and their corresponding colors are denoted on the top. Right: IMC-based estimates of acinar, ADM, and tumor cell percentages using cell segmentation and color intensity quantitation.

We identified significantly highly expressed genes specific to ADM cells (**Figure 4F**). We did this by first performing DEG analysis between ADM and acinar cells, ADM and tumor cells, and tumor and acinar cells. Then, from the ADM comparisons, we removed the DEGs between tumor and acinar cells which are largely driven by acinar-ductal cell type-specific differences. Intriguingly, the most highly expressed ADM genes were oncogenic drivers, including *CD81, FXYD2, SPP1, IGFBP7, VIM*, and *MGP*, as well as tumor suppressors, *A2M, RGS5, TIMP3,* and *SPARCL1*. One possible explanation of this phenomenon is that tumor suppressor genes may be upregulated in response to oncogene expression in this transitional state, evident by their increased expression in the ADM Mutated group, which represents later stage ADM. During the ADM transition from acinar to ductal or ductal to acinar, it may be that these upregulated features are a result of the transformation process towards either cell fate. We observed that *VIM*, an epithelial-to-mesenchymal transition (EMT) marker, is significantly overexpressed in ADM and ADM Mutated (**Figure S4A).** We then analyzed the expression of other EMT-related and stem cell genes to determine whether these pathways were also associated with dedifferentiation of acinar cells into ductal cells (**Figure 4G**). Indeed, compared to acinar cells, both tumor and ADM cells highly express EMT and stemness genes. Interestingly, when comparing ADM and tumor cells, we found a largely mutually exclusive pattern of expression between a subset of genes in each pathway, particularly EMT genes, which are highly upregulated in the ADM Mutated population, and to a lesser extent, the ADM population as well (**Figure 4G**). In the EMT-related genes for instance, *MMP3*, *SNAI2*, and *GCLC* are highly expressed in tumor cells only, while *MMP2*, *VIM*, and *ITGB3* are more highly expressed in both ADM groups, particularly in the ADM Mutated cells.

While we detect low ADM cells in samples throughout these two cases, a vast majority of ADM cells were captured from one sample in HT122P1 (H3), one sample in HT168P1 (H3) and the NAT sample from HT168P1. In HT122P1, sample H3 is the only spatial sample with significant acinar and ADM fractions (**Figure 4H**). To further characterize the ADM heterogeneity in a manner that preserves spatial integrity, we used imaging mass cytometry (IMC) enlisting anti-pan-keratin to label tumor cells and anti-α-amylase to detect acinar cells (**Figure 4I, STAR Methods**). This revealed differences in the dominant tumor morphology between samples H3 and H9, with a patch-like, poorly differentiated morphology in H3 with intermixed acinar cells and a more ductal-like, well-differentiated morphology in H9, H4, and H5 (**Figures 4I and S4B-C**). In order to approximate the amounts of tumor, ADM, and acinar cells from the IMC images, we used a cell segmentation approach and quantified the intensity of α-amylase and pan-keratin for each cell (**STAR Methods**). Using a heuristic in which the presence of both labels indicates ADM cells, we quantified the numbers of tumor, ADM, and acinar cells. This result was remarkably consistent with the scRNA estimates (**Figure 4I**).

### Cancer Associated Fibroblasts Subtypes

The TME plays a critical role in promoting tumorigenesis (Uzunparmak et al., 2019; Ren et al., 2018). In PDAC, there are three major subtypes of CAFs: myCAFs, iCAFs, and apCAFs (Elyada et al., 2019; Sahai et al., 2020; Helms et al., 2020). Consistent with the literature, we identified clusters of iCAFs, myCAFs, and apCAFs. We also observed two subpopulations of iCAFs with unique expression profiles, which we denote as CXCR4+ iCAFs and CD133+ iCAFs (**Figures 5A and 5B**). Several CAF markers such as *ACTA2* and *FAP* are commonly used to identify CAF subtypes; however, they are not definitive markers and are often expressed in both iCAFs and myCAFs (Kraman et al., 2010). We observed that *TAGLN* and *ACTA2* discern myCAFs, while *FAP* and *CXCL12* distinguish iCAFs (**Figure 5B**). apCAFs were identified by expression of *HLA-DRA* and *CD74*, among others (**Figures 5B and S5A**). CXCR4+ iCAFs and CD133+ iCAFs are defined by very high expression of *CXCR4* and CD133 (*PROM1*), respectively, although they also weakly express myCAF and apCAF marker genes (**Figure 5B**). We observed that while most CAFs in every specimen of every patient tumor are iCAFs or myCAFs, the other CAF subtypes are present at low numbers throughout (**Figure 5C**). Notably, the CD133+ iCAFs comprise a large proportion of CAFs in one gemcitabine+nab-paclitaxel tumor (HT122P1) but were only recovered from two out of the four spatial samples (**Figures 5C and S5B**). These CD133+ iCAFs express several cancer stem cell gene markers, including *CD133, MET, EPCAM, CD24,* and *CD44*, with some genes expressed at an even higher level than the tumor cells themselves (**Figure 5D**). Interestingly, we observed high *CD44* expression in apCAFs and CXCR4+ iCAFs as well. Furthermore, *VIM* and *NFE2L2* were most highly expressed in apCAFs, which were more abundant in treated samples (p < 10^−5^) (**Figure 5E**). These results suggest that small unique CAF subpopulations that express cancer-driving programs exist within standard CAF subtypes.

**Figure 5:**
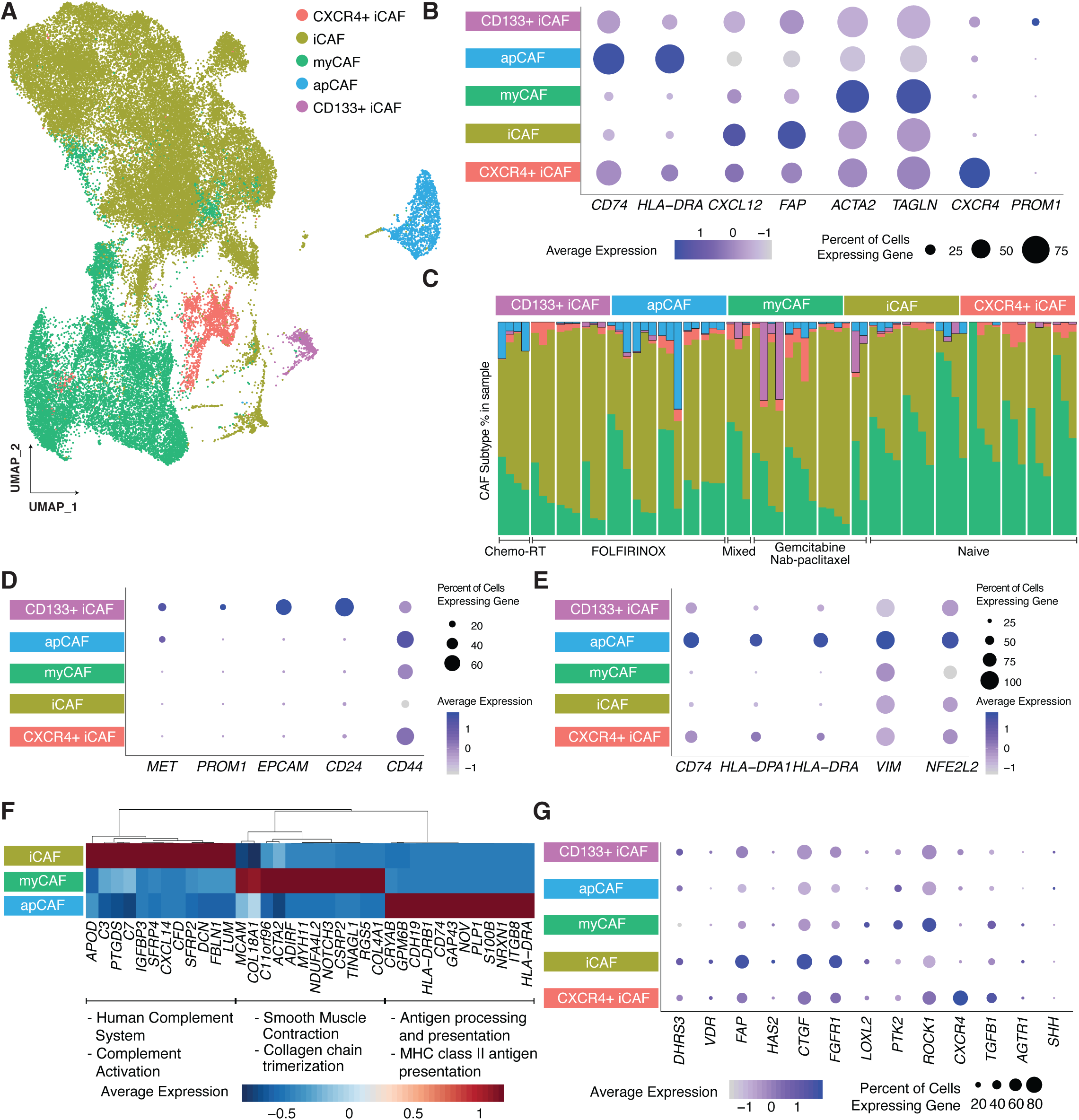
Cancer-Associated Fibroblast Subtypes. A) UMAP of CAF subtypes. **B)** A subset of gene markers used to distinguish between CAF subtypes. **C)** CAF subtype distribution in the cohort and across treatment groups. **D)** Cancer stem cell gene marker expression across CAF subtypes. **E)** Expression of apCAF markers, *VIM*, and *NFE2L2* across CAF subtypes. **F)** Top DEGs across iCAFs, myCAFs, and apCAFs. **G)** Expression of genes currently targeted by clinical trials across CAF subtypes.

To better understand the role of CAFs in tumorigenesis, we analyzed the expression patterns of CAF subtypes to test whether they were enriched for TME-remodeling pathways. Loss of *CAV1* and *CAV2* are associated with the CAF phenotype, which is associated with poor clinical outcomes (Chatterjee et al., 2015; Chen & Che, 2014). Aside from myCAFs, we observe a significant reduction of *CAV1* and *CAV2* expression in all CAFs compared to fibroblasts present in NAT samples (**Figure S5C**). This suggests that myCAFs may have a different biological role in the tumor microenvironment. In addition to their very high levels of CXCR4 expression, CXCR4+ iCAFs express very high levels of its ligand, *CXCL12* (**Figure S5D**) (Liekens et al., 2010; Peitzsch et al., 2015; Eckert et al., 2018). Also, while apCAFs have high expression of *NFE2L2,* which is involved in oxidative damage repair, myCAFs have high expression of *HIF1A,* which regulates tolerance to hypoxic environments (**Figure S5E**). Together, this suggests that different CAF subtypes may play different roles in remodeling of the TME (Semenza, 2003; Huang & Taniguchi, 2017).

We identified the top DEGs between our main CAF subtypes (apCAF, iCAF, and myCAF) (**Figure 5F**). myCAFs upregulate genes that are part of the smooth muscle contraction and collagen chain trimerization pathways, including *MCAM*, *ACTA2*, and *NOTCH3*. iCAFs upregulate genes that are part of the complement system and complement activation pathways, including *IGFBP3*, *PTGDS*, and *CXCL14*. apCAFs upregulate genes that are part of antigen presenting pathways, including *MCAM*, *ACTA2*, and *NOTCH3*. Additionally, we compared the expression of CAF genes currently targeted by clinical trials registered as of 01/2020 (Sahai et al., 2020) (**Figure 5G**). As treated samples compared to untreated samples have a depletion of myCAFs and enrichment of iCAFs, the effectiveness of additional therapies targeting CAFs may differ across treatment groups (**Figure 5C**).

### Immune Populations and Their Interaction with Tumor Cells

To obtain more insight and identify potential ways to address the immunosuppressed TME characteristic of PDAC (Unzunparmak et al., 2019), we identified and reclustered immune cells into two major classes: lymphocytes or myeloid and dendritic cells. In the latter group, we further distinguish between type I and type II classical dendritic cells (cDC1, cD2), macrophages, monocytes, and neutrophils (**Figures 6A, S6A, and S6B**). We observed that myeloid cells and cDCs strongly express TME remodeling pathway genes, such as angiogenesis and hypoxia pathways, including *TGFB1, NFE2L2 (*Nrf2*), VEGFA,* and *HIF1A* at higher levels than tumor cells (**Figure 6B**). Furthermore, we observed high expression levels of genes within the Nrf2 pathway, which regulates oxidative damage repair, including *NQO1* and *GPX2* (**Figure 6C**). While tumor cells do not have significant expression of *NFE2L2*, it does have activation of the pathway. Therefore, such activation may be triggered via paracrine interactions with TME cells and would indicate that myeloid cells and dendritic cells contribute towards a pro-tumor TME.

**Figure 6:**
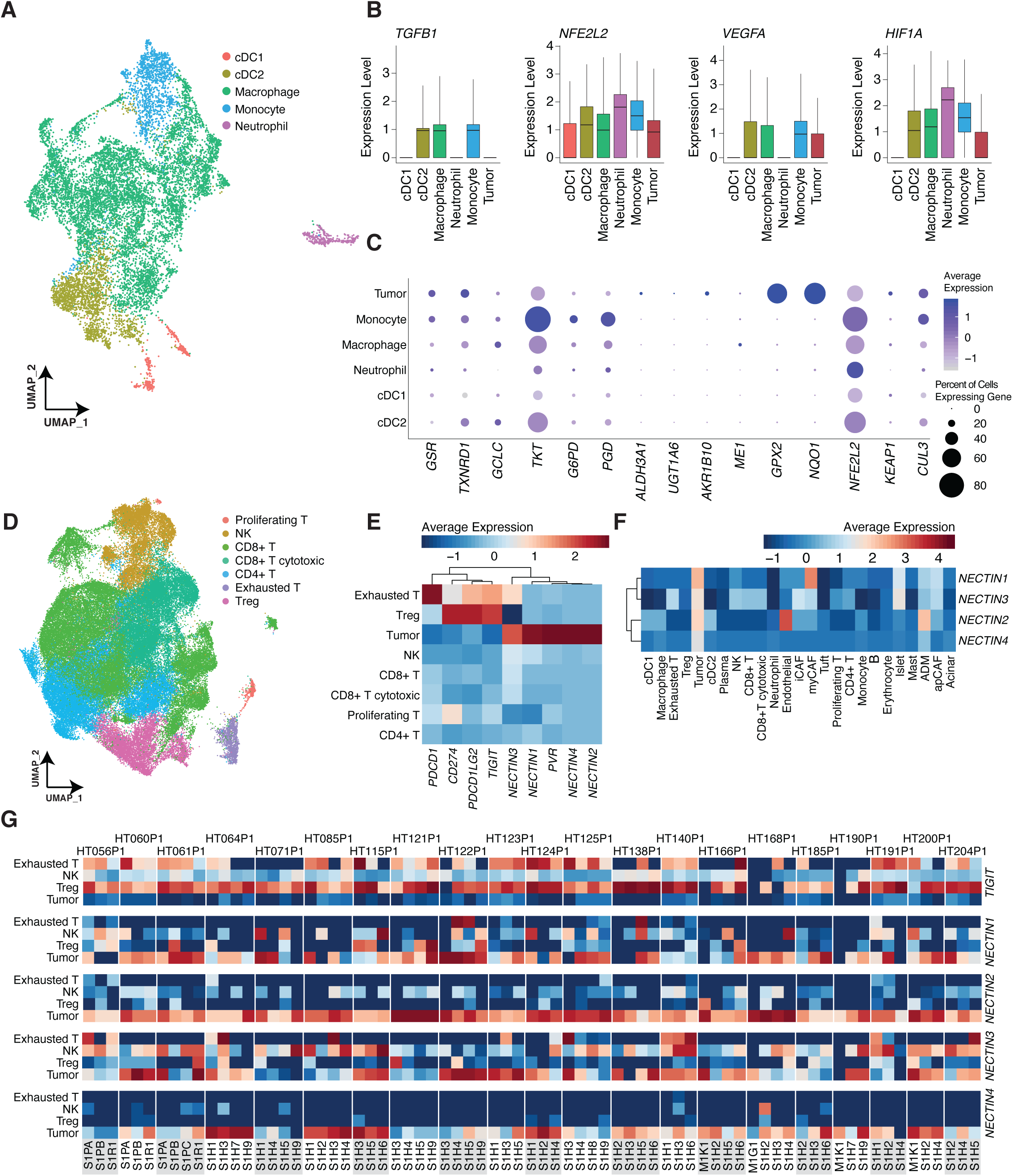
Immune Interactions in the Tumor Microenvironment. A) Myeloid cells and dendritic cell clusters. **B)** *TGFB1, NFE2L2, VEGFA,* and *HIF1A* expression across cell types. **C)** Nrf2 pathway gene expression. Tumor cells are included for comparison. **D)** Lymphocyte and NK cell clusters. **E)** *TIGIT* and nectin receptors expression in tumor, NK, and lymphocyte cells. PD1, PD-L1, and PD-L2 are included for reference. **F)** Expression of nectin receptors across all cell types. **G)** Average expression of *TIGIT, NECTIN1, NECTIN2, NECTIN3,* and *NECTIN4* in exhausted T cells, NK cells, Tregs, and tumor cells.

Within the lymphocyte and NK subset, we assigned states to CD4+ and CD8+ T based on their exhaustion, proliferation, and cytotoxic markers (**Figures 6D, S6A, and S6B**). We observe similar percentages of cell types across treatment groups, with slightly higher abundance of CD4+ T cells in FOLFIRINOX samples and higher numbers of CD8+ T cells in treated samples (**Figure S6C**). CD4+ T cells and Tregs in FOLFIRINOX samples had high expression of heat shock genes, such as *HSPA1A, HSPA1B, HSPH1,* and *HSPD1,* compared to other treatment groups. (**Figure S6D**). Further, pathway enrichment analyses revealed a large number of cellular responses to heat stress in both CD4+ T cells and Tregs in these samples (**Figure S6E**). Receptor-ligand analyses furthermore reveal an interaction between the *TIGIT* receptor in lymphocytes and its *NECTIN2* receptor across all samples, which we found is highly expressed in tumor cells (**STAR Methods**). This is consistent with a previous report that *NECTIN4* has high tumor specificity and is a potential target for immune checkpoint blockade (Reches et al., 2020; Gorvel and Olive, 2020). *TIGIT* interaction with NECTIN receptors inactivates T cell and NK function, which could be used by the tumor for immune evasion. We expanded our analysis to all nectin receptors and observed that *NECTIN1*, *NECTIN2*, and *NECTIN4* are all expressed solely in tumor cells, but *NECTIN3* is expressed in some lymphoid cell types, while *TIGIT* is largely expressed in Tregs and exhausted CD4+ T cells (**Figure 6E**). This suggests that this interaction may be contributing towards the immunosuppressive TME in PDAC. Furthermore, we observe that *PD-L1* and *PD-L2* are not expressed in tumor cells at all, consistent with the poor response of PDAC to anti-*PD-1/PD-L1* immunotherapy (Feng et al., 2017; Birnbaum et al., 2016; Pu et al., 2019). We extend our analysis of nectin receptors to all cell types to assess tumor-specific expression beyond lymphocytes and note that while nectins are overall quite tumor-specific, *NECTIN1* is highly expressed in myCAFs, *NECTIN2* in endothelial cells, and *NECTIN3* in islet cells; *NECTIN4* is the most tumor cell-specific NECTIN receptor (**Figure 6F**), consistent with previous reports (Reches et al., 2020). We analyzed TIGIT and Nectin receptors expression at individual sample levels in Tregs, NK cells, exhausted T cells, and tumor cells (**Figure 6G**). Consistent with the previous analysis, we observed high expression of all nectin receptors in tumor cells and *TIGIT* in Tregs and exhausted T cells, but also noted a substantial degree of heterogeneity across cases, particularly in *TIGIT* expression in exhausted T cells and in *NECTIN1* and *NECTIN3* expression in tumor cells. These results suggest a rationale for targeting the TIGIT-NECTIN axis in PDAC.

### Cell Type Biomarkers and Treatment Implications

Bulk RNA-Sequencing data of tumors reflects gene expression signatures originating from a mixture of cell types. Unless the samples have exceptionally high tumor purity, bulk analysis cannot be a completely faithful representation of tumor cell gene expression (Nieuwenhuis et al., 2020). scRNA Seq data enables the identification of gene expression patterns from specific cell type populations. By analyzing only the tumor cells in the scRNA data, we identified 109 tumor markers, 56 of which are expressed on the cell surface, that are highly expressed in tumor cells compared to all other cell types, with a small subset of markers (n = 15) also strongly expressed in ADM cells (**Figure 7A, STAR Methods, Table S3**). Genes that are highly expressed in both tumor cells and ADM include *MMP7, CXCL17, TFF1, LGALS4,* and *CEACAM5*. Additionally, within the 109 genes, we found 25 genes that are significantly differentially expressed between tumors treated with different treatment regimens (**Figure 7B**). We identified several specific genes and proteins that are significantly highly expressed at the bulk RNA and protein levels in the gemcitabine+nab-paclitaxel, treatment-naïve, and FOLFIRINOX groups (FDR < 0.05) (**Figure 7B**). Notably, while *KRT17* and *C19orf33* are commonly used as markers to distinguish between epithelial and neoplastic cells, we observed large differences in expression in FOLFIRINOX-treated samples compared to other samples. We also observe a set of genes that overlap of our previously identified markers for the single cell-based classical subtype. These genes include *TFF1, TFF3, OLFM4,* and *CLDN18*, which suggest that in addition to treatment status, tumor subtypes also have an impact on specific tumor biomarker expression (**Figure 7B**).

**Figure 7:**
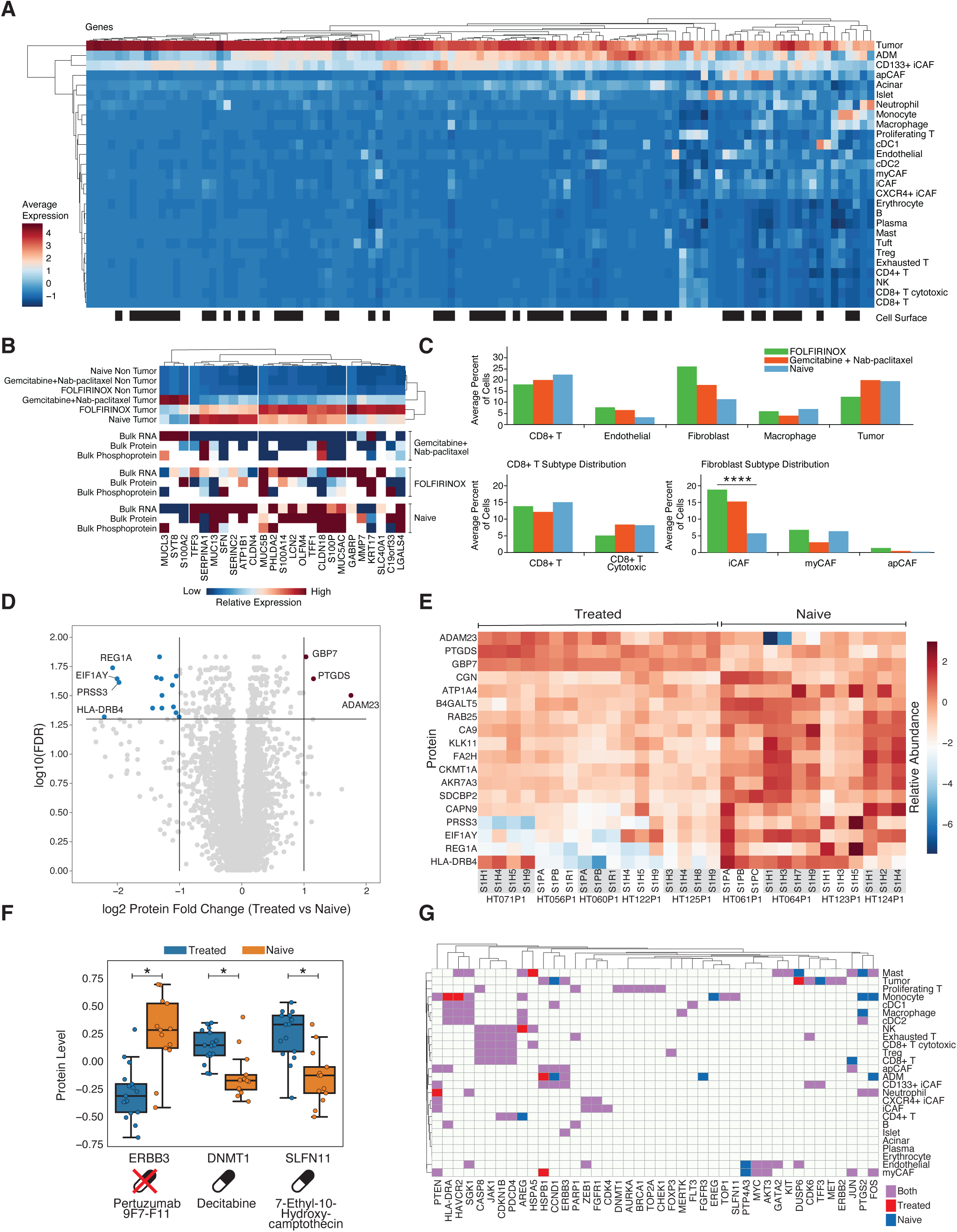
Tumor Biomarkers and Druggable Targets. A) Average expression of tumor biomarkers across all cell types. **B)** Top: Significantly differentially expressed genes by treatment groups and tissue status. Lower panels: scRNA, bulk RNA, proteomics, and phosphoproteomics levels of tumor biomarkers that are differentially expressed across treatment groups. **C)** Top: cell type percentages split by treatment groups. Bottom: cell type distributions for CD8+ T cells (cytotoxic and no particular substrate) and CAF subtype (iCAF, myCAF, and apCAF). **D)** Top differentially expressed proteins across treated and untreated samples. **E)** Protein levels of 18 differentially expressed proteins across all samples with proteomics data. **F)** Protein levels of ERBB3, DNMT1, and SLFN11 in treated and untreated samples. **G)** Druggable genes significantly upregulated across cell types.

In order to determine the differences that occur following treatment, we assessed changes in cell type proportions within each of the three major treatment groups (**Figure 7C**). The largest difference was in fibroblasts, where both treated groups had higher numbers of fibroblasts than the treatment naive group. Notably, we determined that the difference is driven by a 3-fold higher amount of iCAFs in FOLFIRINOX and Gemcitabine+Nab-paclitaxel samples (p < 10^−4^), with little difference in myCAF abundance between treatment groups. As iCAFs are understood to be pro-tumorigenic (Hosein et al., 2020), this large increase of iCAFs aftertreatment may bode poorly in terms of treatment resistance. While we detected a decrease in CD8+ T cells, particularly cytotoxic T cells, and tumor cells in FOLFIRINOX samples, these differences were not significant. These observations provide evidence that treated tumors have much higher levels of iCAFs in particular, and represent an attractive potential target for chemoresistant tumors.

We further analyzed the DEGs between treatment groups within each cell type (**STAR Methods**). Acinar cells and endothelial cells harbored the strongest differences in expression between treated and untreated samples. In acinar cells, we detected high expression of regenerating family member genes, such as *REG1B* and *REG3A* in the naïve group only. Higher expression of amylases and lipase genes were found in both treated groups, which have been previously associated with chronic pancreatitis and pancreatic cancer (**Figure S7A**) (Raphael et al., 2016; Stotz et al., 2020; Lasher et al., 2019). In endothelial cells, we observed an upregulation in metallothioneins in treated samples, particularly in gemcitabine+nab-paclitaxel samples. (**Figure S7B**). Interestingly, *PRSS1* and *PRSS2*, which are associated with hereditary pancreatitis, are highly expressed in FOLFIRINOX endothelial cells only (Le Maréchal et al., 2006; Whitcomb et al., 2012). Similar to acinar cells, we observed high expression of lipase, and chymotrypsin genes in endothelial cells but only in FOLFIRINOX-treated cases.

With respect to the tumor cells, we identified DEGs between treated and treatment-naïve samples. The top DEGs include *MMP7*, *REG4*, and *VCAN* in treated tumor cells and *TFF3, ITGB8, and PLCG2* in treatment-naïve tumor cells (**Figure S7C**). Breaking down the treated group into FOLFIRINOX and gemcitabine+nab-paclitaxel cells results in the same top DEGs in the naïve group, but we detect that *VCAN* and *MMP1* to be gemcitabine+nab-paclitaxel-unique (**Figure S7D**). We applied a similar comparison to the bulk RNA and protein data, which reflect a mixture of cell types and once again compared treated vs treatment-naïve samples (**STAR Methods**). The genes with the greatest fold change between treated and untreated samples included *AMY2A, REG1B,* and *CTRB2* in the untreated samples and *KRT5* and *KRT6A* in the treated group. The first set of genes are known to be acinar gene markers and the second set is composed of epithelial genes, suggesting that most differences are driven by cell-type differences due to low tumor purity (**Figure S7E**). While we do not observe significant differences in the tumor or acinar cell proportions estimated from the paired scRNA data, the tissue portion subjected to bulk sequencing was not completely identical to the scRNA tissue, and it is possible that this skewed the observed results (**STAR Methods**). At the protein level, we did not observe cell-type differences. Focusing on proteins with a 2-fold or greater change, we identified 18 differentially expressed proteins (DEPs) between treated and untreated samples and find that GBP6, PTGDS, and ADAM23 are elevated in treated samples while REG1A, EIF1AY, PRSS3, and HLA-DRB4 are elevated in naïve samples, among others (**Figure 7D**).

While these 18 proteins overall display such patterns between treated and naïve samples, we observe a modest amount of heterogeneity between spatial samples and between a subset of tumor cases (**Figure 7E**). For instance, treated tumor HT071P1 has high levels of HLA-DR4 while treated tumor HT122P1 has high levels of EIF1AY. Similarly, naïve tumor HT064P1 has very low levels of ADAM23 only in 2/4 spatial samples, suggesting spatial heterogeneity. In order to determine whether this heterogeneity is due to differences in the composition of non-tumor cell types, we assessed whether any of these 18 proteins were significantly differentially expressed between treatment groups in specific cell types in the scRNA data (**STAR Methods**). Curiously, we found that *REG1A* is upregulated in naïve apCAFs and FOLFIRINOX endothelial cells. Additionally, we observe *PTGDS* overexpression in gemcitabine+nab-paclitaxel iCAFs and *PRSS3* overexpression in acinar cells in both treated groups compared to the naïve cells (**Figure S7F**). These results suggest that several of these differentially abundant proteins may be driven by the TME rather than the tumor cells. Finally, we match all significant DEPs (n = 143, no minimum fold change of 2 filter) with the CiVIC druggable database (Griffith et al., 2017) to identify potential druggable proteins that are differentially expressed in treated and untreated samples (**STAR Methods**). We identified 3 matches to expression-based druggable targets: ERBB3 - Pertuzumab/9F7-F11 in naïve samples and DNMT1-Decitabine and SLFN11-7-Ethyl-10-Hydroxycamptothecin in treatment-resistant samples (**Figure 7F**). These results suggest that DNMT1 and SLFN11, which are known to drive other cancers and are currently targetable, could represent drivers in the resistant tumors and thus may warrant pre-clinical assessment for treatment efficacy. On the other hand, targeting ERBB3 may not be possible in resistant tumors compared to naïve tumors due to treatment-induced changes in protein levels (Murai et al., 2019; Zhang & Xu, 2017).

Lastly, we extended our druggable target analysis to all cell types in the scRNA data. For each cell type, we split the analysis into treated and untreated cells and identified expression-based druggable targets (**Figure 7G**). Most druggable genes identified were present in both treated and untreated cells (i.e., significantly highly expressed in the given cell type but not differentially expressed between treatment groups). By clustering the data, we identified modules of druggable targets that roughly correspond to lymphocytes, myeloid cells, and CAFs. When clustering the drugs that target these genes, these cell type modules are also present (**Figure S7G**). These include *PDCD4, CDKN1B, CASP8*, and *JAK1* in lymphocytes and *HLA-DRA, SGK1*, and *HAVCR2* in dendritic and myeloid cells. Interestingly, apCAFs, ADM cells, and CD133+ CAFs group together with druggable protein products of *HSPB1, CCND1, and ERBB3* and endothelial cells and myCAFs group together with druggable products of *PTP4A3*, *MYC*, and AKT3. We note that only 2 out of 8 tumor targets, *MET* and *ERBB2*, seem to be tumor-specific, implying a potentially lower chance of off-target effects in other cell types.

## DISCUSSION

We conducted a comprehensive multi-omic spatial characterization of PDAC by integrating bulk sequencing and proteomics/phosphoproteomics, single cell sequencing, and high-resolution cellular imaging technologies. Each of these modalities makes a crucial contribution to the overall picture. Spatial sampling and imaging mass cytometry orthogonally reveal tumor heterogeneity. Single cell resolution enables the characterization of tumor cells independent of purity level, thus bypassing what has been a substantial challenge in PDAC omic studies, and powers a detailed evaluation of the TME and cell-cell interactions (Raphael et al., 2017). Different treatment groups allowed us to identify potential mechanisms of treatment resistance and identify new targets worthy of further mechanistic inquiry. Over 50% of spatial samples showed high degrees of cell type heterogeneity, which was echoed by pathological review of H&E slides and subtypes from bulk sequencing data. In certain cases, pathologic review assigned different tumor grades to different samples from the same patient, consistent with the molecular heterogeneity observed in omic analyses.

We identified ADM cell populations, which express both oncogenic features and tumor suppressor genes, and significantly upregulate EMT and stem cell genes, compared to tumor cells (Xu et al., 2019). While the ADM state is important in post-injury pancreas regeneration, it can also transition through intraepithelial neoplasia lesions to full-blown PDAC (Storz, 2017). The unique expression pattern of ADM as an intermediate state suggests there may be a dynamic transition between tumor and acinar fates in ADM, which may lead to the progression towards PDAC when the oncogenic side prevails, commonly due to the acquisition of a driver *KRAS* variant. This would be consistent with the sensitivity of acinar cells to *KRAS* mutations as a catalyst for ADM and inclination toward PDAC (Xu et al., 2019). It would also further connect our observations of KRAS mutated clusters upregulating *REG1A* and elevation of REG1A in treatment-naïve samples, as REG1A has been proposed as a diagnostic and prognostic marker for PDAC (Li et al., 2016). We also observed spatial heterogeneity in ADM populations, which was associated with more poorly differentiated tumor phenotype.

The function of cancer-associated fibroblasts (CAFs) has also been poorly understood in PDAC.Historically presumed to be staunch drivers of cancer, they are now known to have a much more complicated, even dual behavior that can either drive or suppress cancer development, depending upon numerous factors (Gieniec et al., 2019; Pereira et al., 2019). Indeed, their highly heterogeneous nature has stimulated efforts to discover and catalog CAF types. Here, we identified iCAFs, myCAFs, and apCAFs, further classifying two small iCAF subsets as CD133+ and CXCR4+. The CD133+ iCAFs are characterized by high expression of cancer stemness genes, including *PROM1 (CD133), MET, EPCAM,* and *CD24.* We identified markers and activated pathways in these subtypes, determining that most CAF genes in current clinical trials are differentially expressed between subtypes and therefore may respond differently (Sahai et al., 2020). Importantly, we observed a distinct pattern of higher iCAF abundance in treated samples. This is important to consider, as IL-1-mediated signaling and JAK-STAT signaling in iCAFs has motivated respective studies of adding IL-1R blockade to standard-of-care (FOLFIRINOX-based) chemotherapy (Hosein et al., 2020; ClinicalTrials.gov: NCT02021422). Additionally, treating KPC mouse models with a JAK inhibitor to target iCAFs has resulted in decreased tumor size (Biffi et al., 2019).

Immunotherapy continues to progress across the cancer treatment realm, but clinical application is often still plagued by side effects stemming from over-stimulation of the immune system. In some instances, these effects can be particularly serious or even fatal (Wang et al., 2018). Searches for ever-greater specificities to alleviate what have become known as “immune-related adverse events” thus continues (Reches et al., 2020). Single cell analysis revealed that the nectin receptors, in particular *NECTIN4*, are tumor-specific. Receptor-ligand analysis uncovered a potential interaction with TIGIT in Tregs and exhausted T cells, which may be inhibiting the activation of T cell and NK functions. This suggests the possibility that the nectin-TIGIT interaction may be a target worth exploring in PDAC. Consistent with previous analysis, we observed high expression of all nectin receptors in tumor cells and *TIGIT* in Tregs and exhausted T cells but also noted a substantial degree of heterogeneity across cases, particularly in *TIGIT* expression in exhausted T cells and in *NECTIN1* and *NECTIN3* expression in tumor cells. The specificity problem in immunotherapy is increasingly urgent for advancing the safe and effective application of inhibitors.

In conclusion, this study provides a comprehensive analysis of PDAC spatial heterogeneity and treatment characteristics by integrating bulk and single cell omics and imaging technologies. We determined high levels of heterogeneity in PDAC including spatially separated driver clones, KRAS mutated and KRAS WT tumor populations, tumor-ADM interactions, and subtype heterogeneity within the same patients. These results underscore both the utility of spatial sampling from multiple regions of the tumor for characterization, as well as the clinical challenge of capturing PDAC heterogeneity in clinical assays. The identification of biomarkers and differentially expressed genes and proteins between treatment groups as well as cell types provides a resource to identify new targets with clinical relevance.

## ACKNOWLEDGEMENTS

We thank the patients, staff, and scientists who contributed to this study as well as the NCI and the HTAN consortium. All HTAN consortium members are named at humantumoratlas.org. We also thank the Siteman Cancer Center and the McDonnell Genome Institute for their support. The following grants supported this work: U2CCA233303 to L.D., R.C.F., W.E.G., and S.T.O.; U24CA211006 to L.D.; U24CA209837 to K.I.S.; R01HG009711 to L.D. and F.C..

## AUTHOR CONTRIBUTIONS

**Study Conception & Design**: L.D., R.C.F., W.E.G., F.C., S.T.O.

**Developed and Performed Experiments or Data Collection**: R.G.J., J.M.H., C.C.F., L.T., E.L.A., S.E.C., X.Y., K.S., A.N.R., J.P., L.C., F.C., B.F., C.F., A.N.R., R.V., M.S., K.J.R., C.O., L.C., S.V.P., D.C., W.G.H.

**Computation & Statistical Analysis**: D.C.Z., R.G.J., E.S., C.M., Y.W., N.D., M.A.W., L.W.,

Y.L., C.Z., R.L., N.V.T., H.Z., H.S., F.W., M.S., H.Z.,

**Data Interpretation & Biological Analysis**: D.C.Z., R.G.J., E.S., J.M.H., C.Z., K.J.R., D.C., M.G.C., D.G.D., W.E.G., R.C.F., L.D., J.F.M., K.C., H.Z., L.Y.

**Writing –** Original Draft: D.C.Z.. Review & Editing: D.C.Z., R.S.F., L.A.F., J.F.M., M.C.W., R.J.M., S.V.P., A.H.K., S.S., K.I.S., T.J., W.G.H., K.C., H.Z., D.C., M.G.C., S.A., D.G.D., S.T.O., F.C., W.E.G., R.C.F., L.D., N.V.T., N.N.

**Administration**: L.D., R.C.F., W.E.G., F.C., S.T.O. S.R.D., E.L.A., L.A.F., C.C.F., R.S.F., J.P., A.H.K., K.I.S., T.J., S.S.

## SUPPLEMENTAL FIGURE LEGENDS

**Figure S1: Cell Types and Sample Similarity, Related to Figure 1. A)** All cells labeled with case ID. **B)** All cells labeled with cell types. **C)** Stroma percentages between treated and untreated samples. **D)** Proteomics PCA. **E)** Phosphoproteomics PCA.

**Figure S2: Bulk Correlations and Single-Cell Subtype Annotations, Related to Figure 2. A)** Stroma score correlations with scRNA estimates. **B)** Immune score correlations with scRNA estimates. **C)** Tumor cells labeled with single-cell subtype. **D)** Tumor cells of HT061P1 labeled with *KRAS* variants. **E)** EMT gene expression in classical and basal-like tumor cells. The overall EMT score is a composite score from 14 EMT related genes.

**Figure S3: Differential Genomic Features in Heterogeneous KRAS Subpopulations, Related to Figure 3. A)** Tumor cell clusters labeled with case ID. **B)** 3 tumor cell clusters originate from most samples, one of which has no mappable *KRAS* mutation. **C)** Top significant DEGs between *KRAS* clusters of multi-tumor origins. **D)** Top significant DEGs between specific *KRAS* hotspot mutations. Only cells with a mappable mutation were included in this analysis. **E)** KRAS and TP53 trans mutation impacts on protein levels. **F)** KRAS and TP53 trans mutation impacts on phosphoprotein levels. Overlapping dots denote several phosphosites from the same phosphoprotein. **G)** *KRAS* mutations in tumor cells of 5 cases with multiple *KRAS* variants mapped. **H)** Tumor cells of HT061P1 labeled with punch of origin. **I)** Arm and gene-level CNV events in HT061P1.

**Figure S4: Additional IMC ROIs for ADM Samples, Related to Figure 4. A)** *VIM* expression. **B)** IMC image of HT122P1 S1H4 sample. **C)** IMC image of HT122P1 S1H5 sample.

**Figure S5: Differentially Expressed Genes in Cancer-Associated Fibroblasts, Related to Figure 5. A)** CAF subtype markers. Tumor cells are included for reference. **B)** Sample and CAF subtype clusters in HT122P1. **C)** *CAV1* and *CAV2* expression. **D)** *CXCR4* and *CXCL12* expression. **E)** *HIF1A* and *NFE2L2* expression. Macrophages and monocytes are included for comparison.

**Figure S6: Immune Population Markers and Heat Shock Gene Expression in FOLFIRINOX Cells, Related to Figure 6. A)** Lymphocyte and NK cell marker expression. **B)** Myeloid and dendritic cell marker expression. **C)** Lymphocyte and NK percentages across treatment groups. **D)** Expression levels of heat shock genes across treatment groups. **E)** Enriched pathways in FOLFIRINOX CD4+ T and Treg populations.

**Figure S7: Differentially Expressed Genes and Druggable Genes in Different Cell Types, Related to Figure 7. A)** Top acinar cell DEGs between treatment groups. **B)** Top endothelial cell DEGs between treatment groups. **C)** Top 5 DEGs each between treated and untreated tumor cells. **D)** Top tumor cell DEGs splitting the treatment group into FOLFIRINOX and Gemcitabine+Nab-paclitaxel groups. **E)** Top bulk RNA DEGs between treated and untreated samples. **F)** scRNA expression of *REG1A*, *PTGDB*, and *PRSS3* in different cell types across treatment groups. **G)** Drugs that target genes significantly upregulated across cell types.

## STAR METHODS

### LEAD CONTACT AND MATERIALS AVAILABILITY

Further information and requests for resources and reagents should be directed to and will be fulfilled by the Lead Contact, Li Ding.

### KEY RESOURCES TABLE

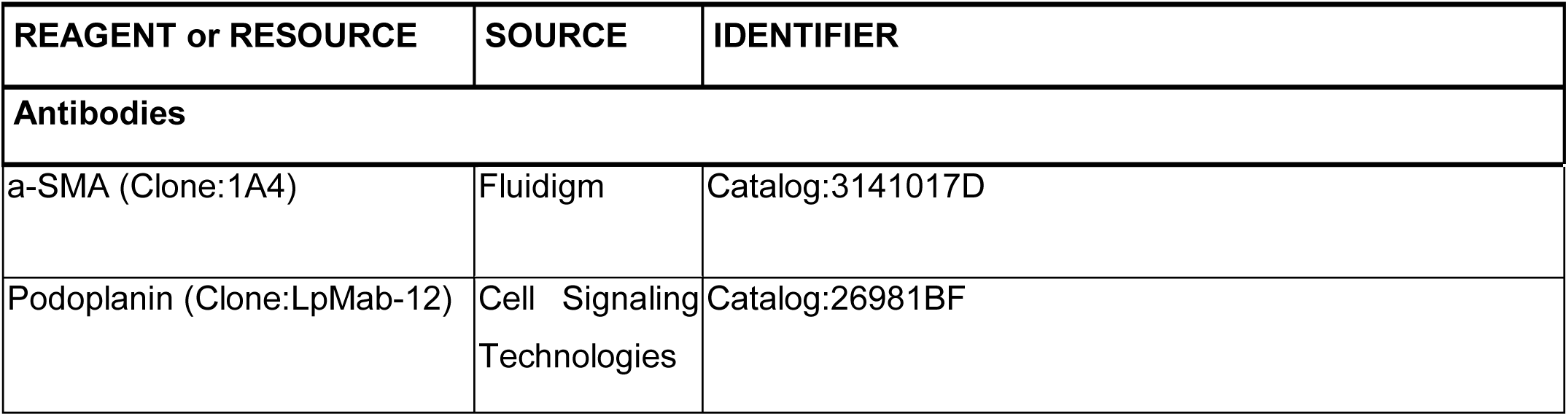
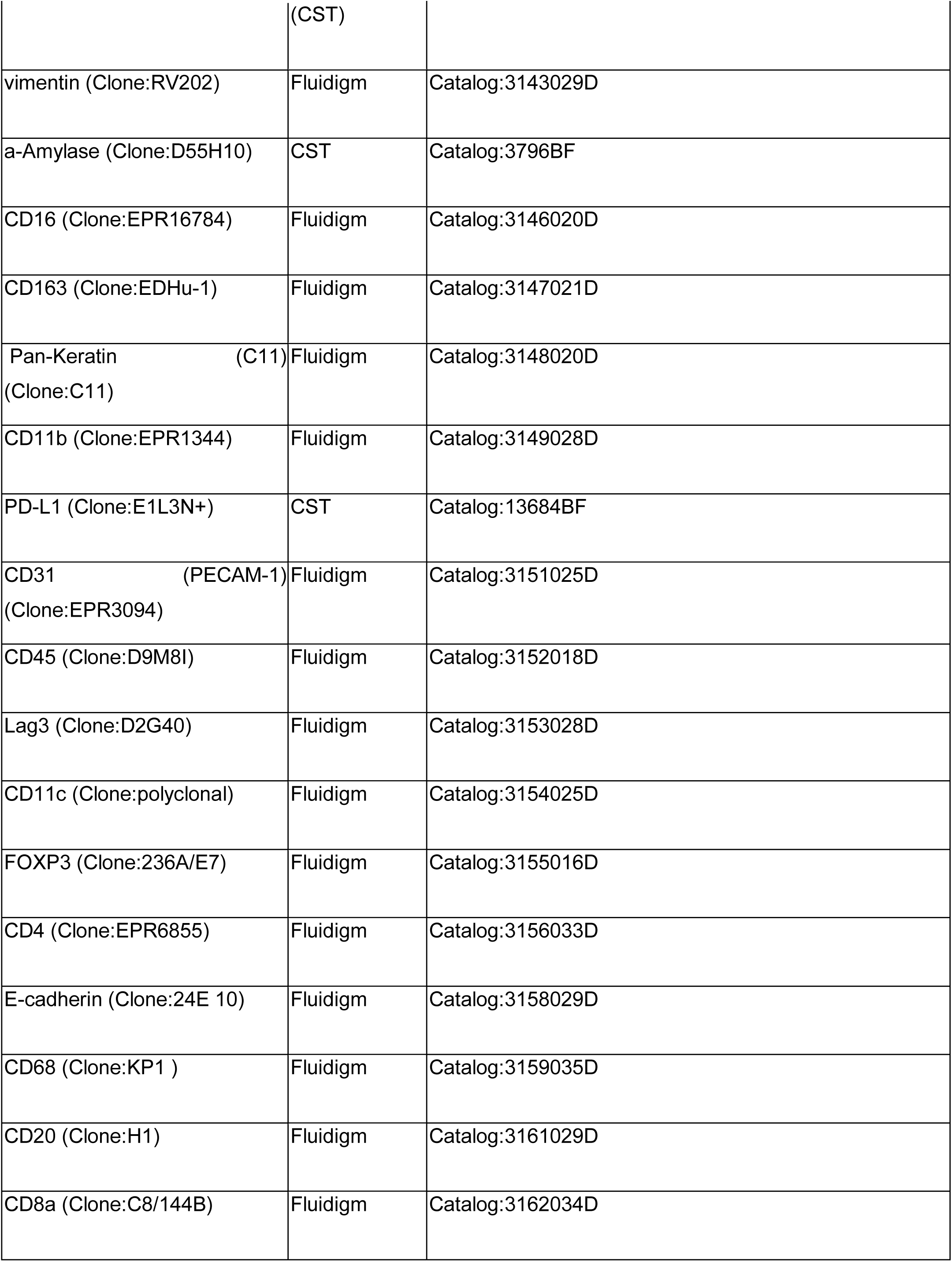
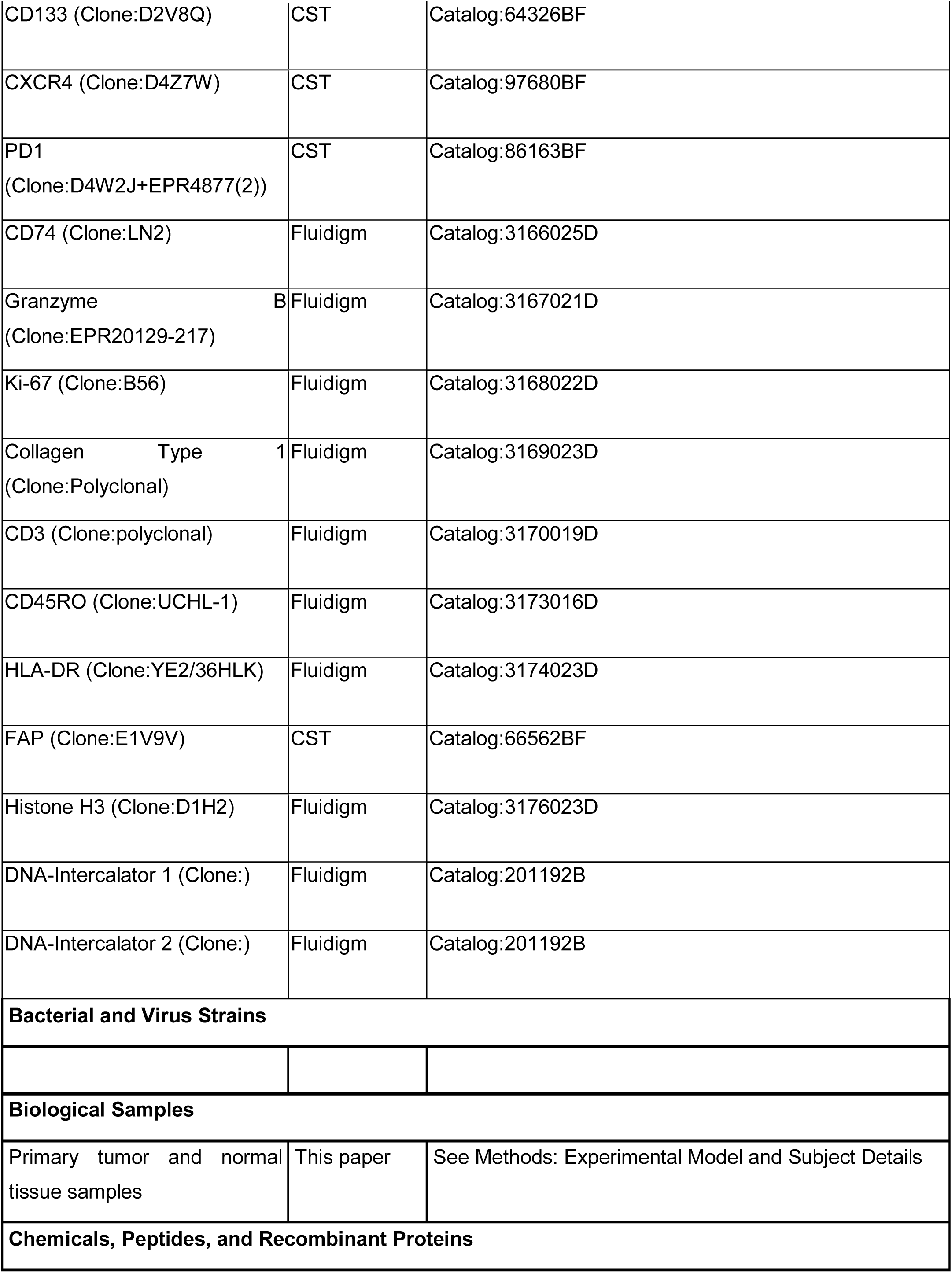
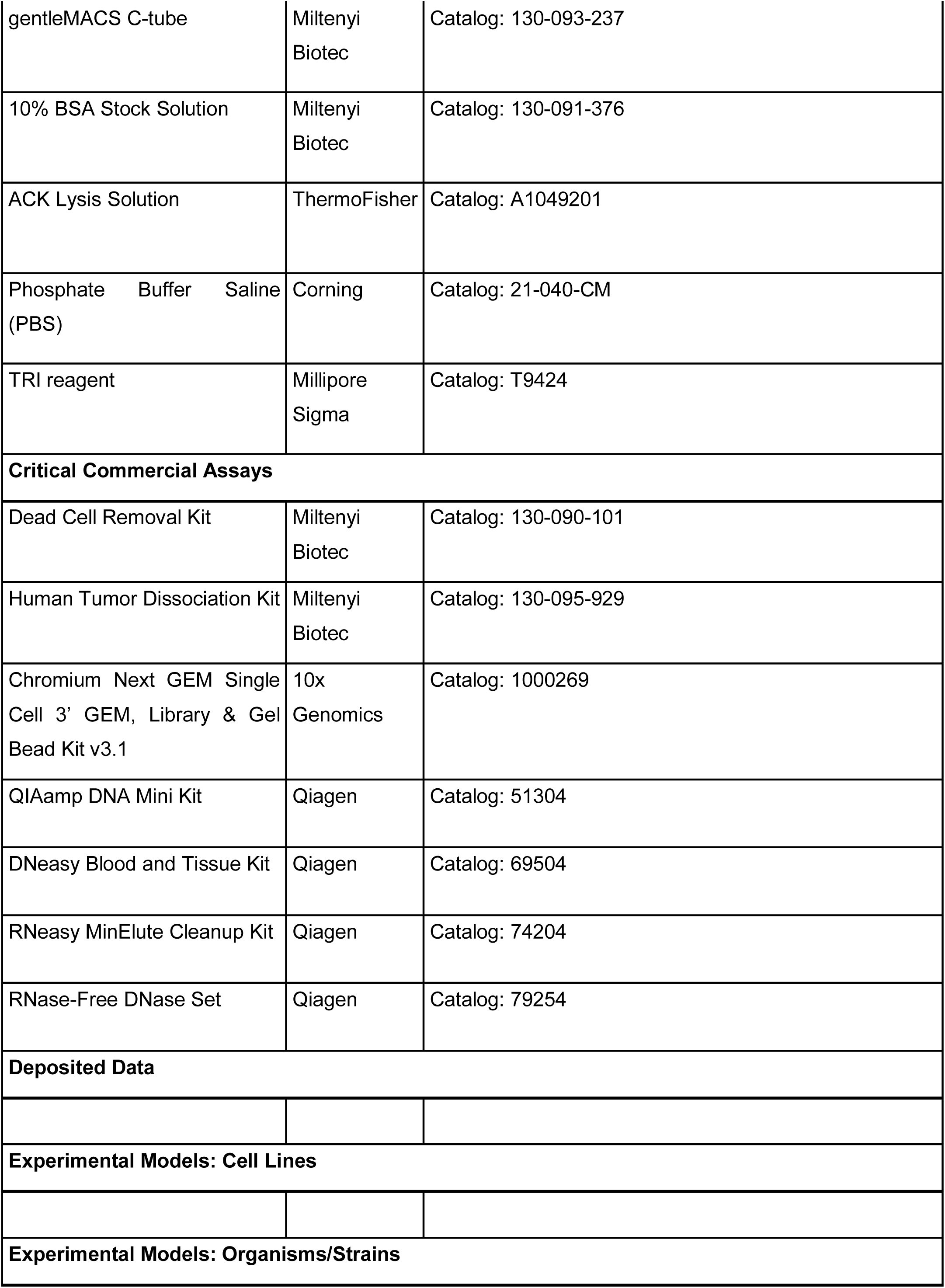
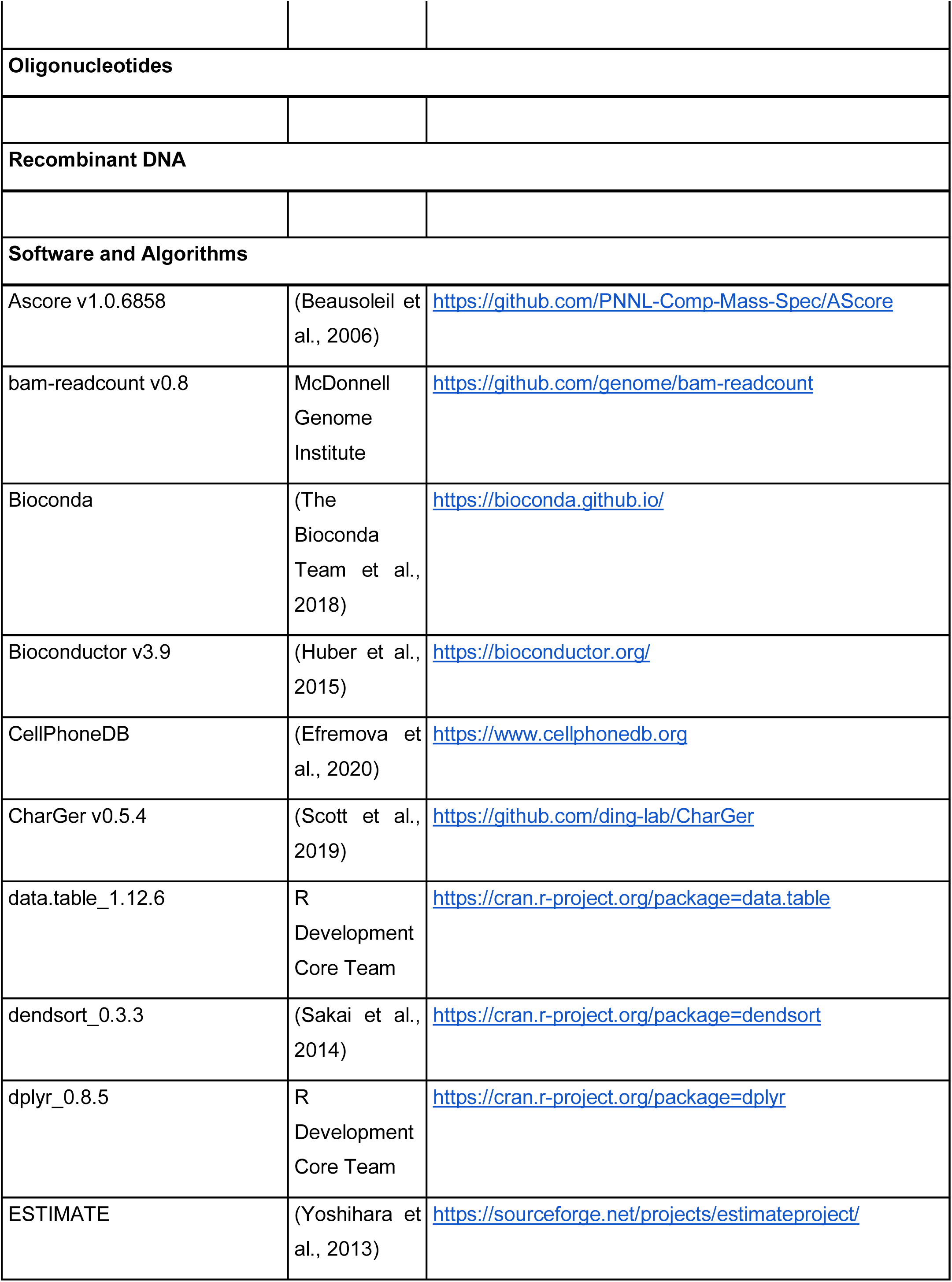
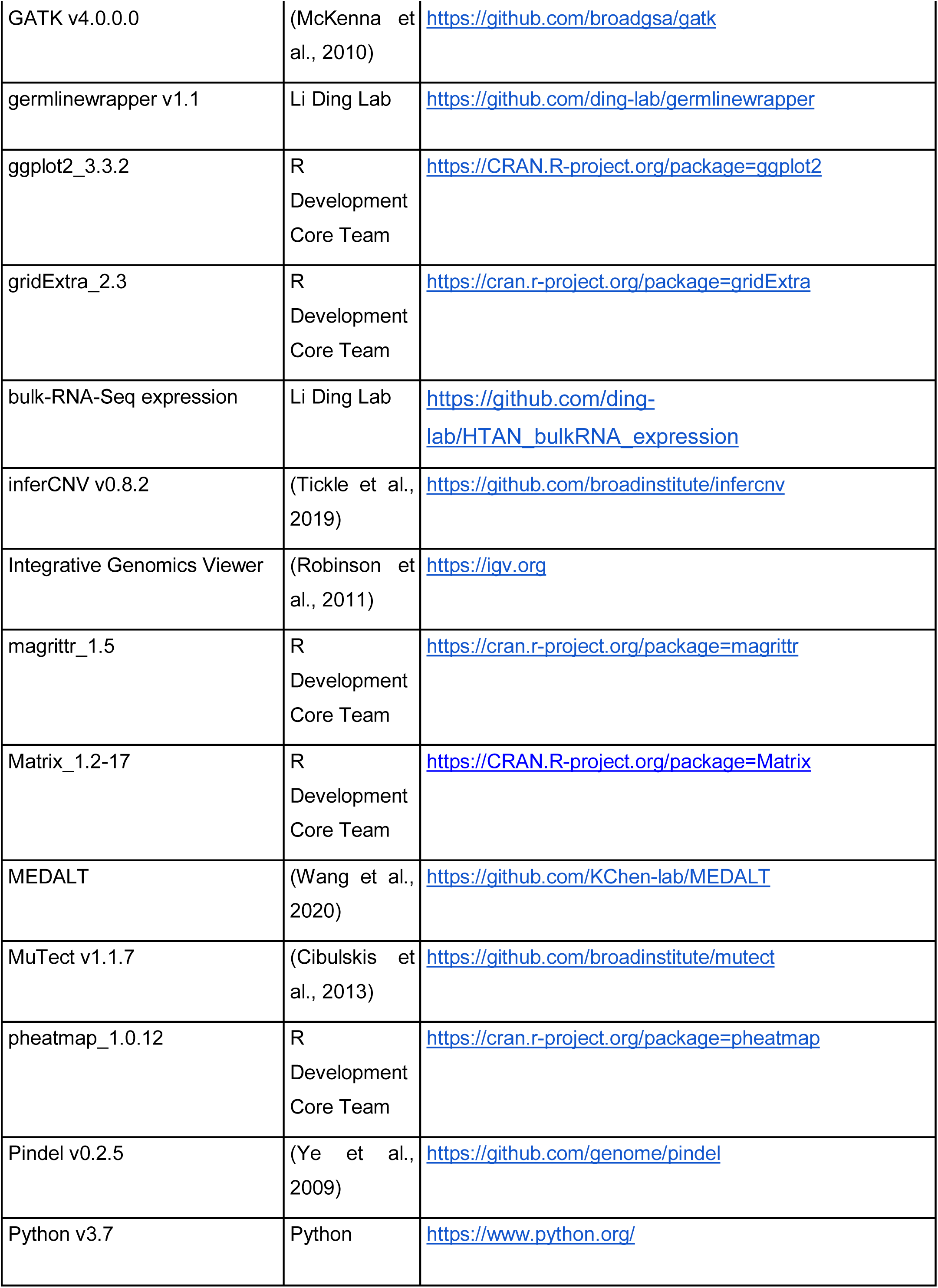
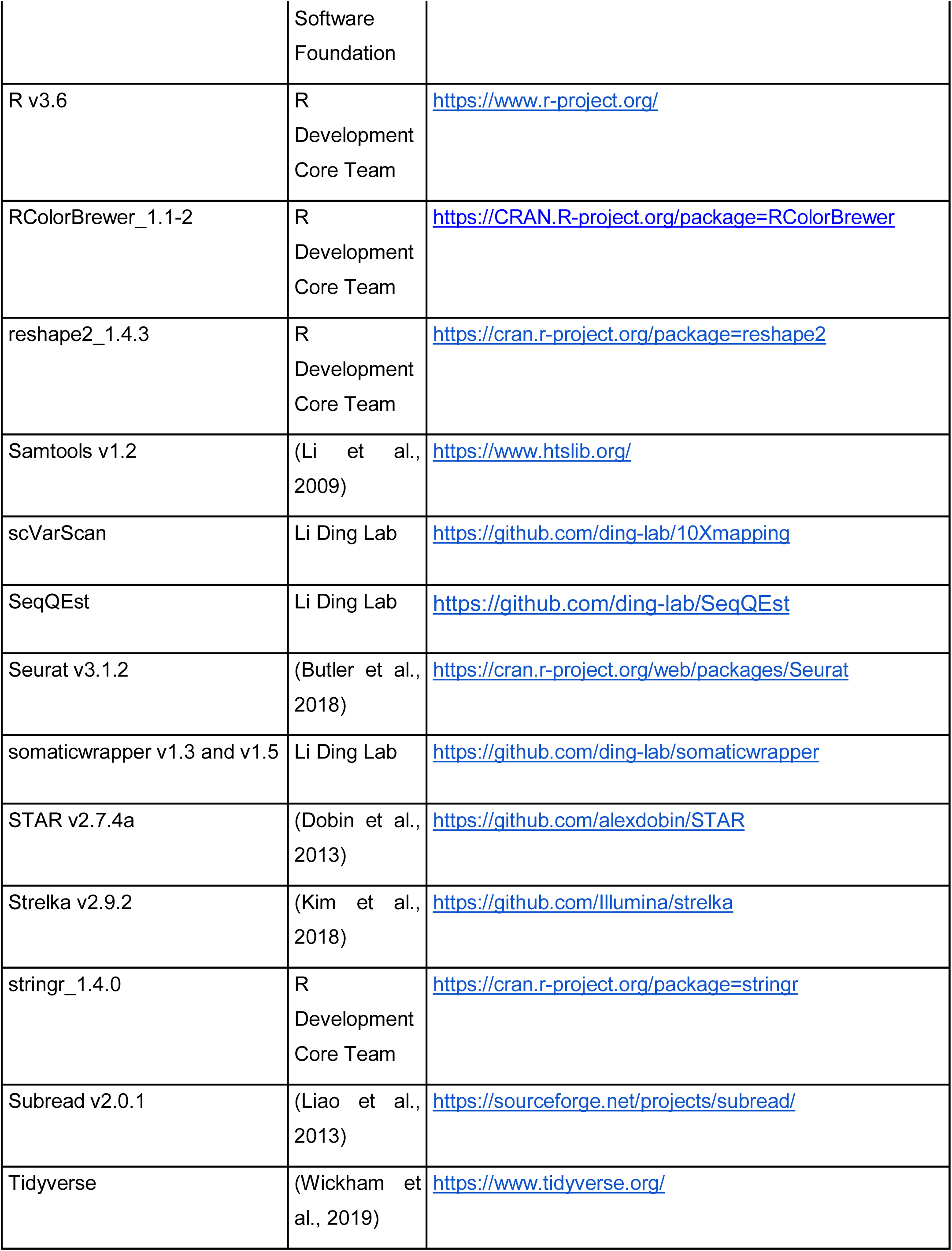
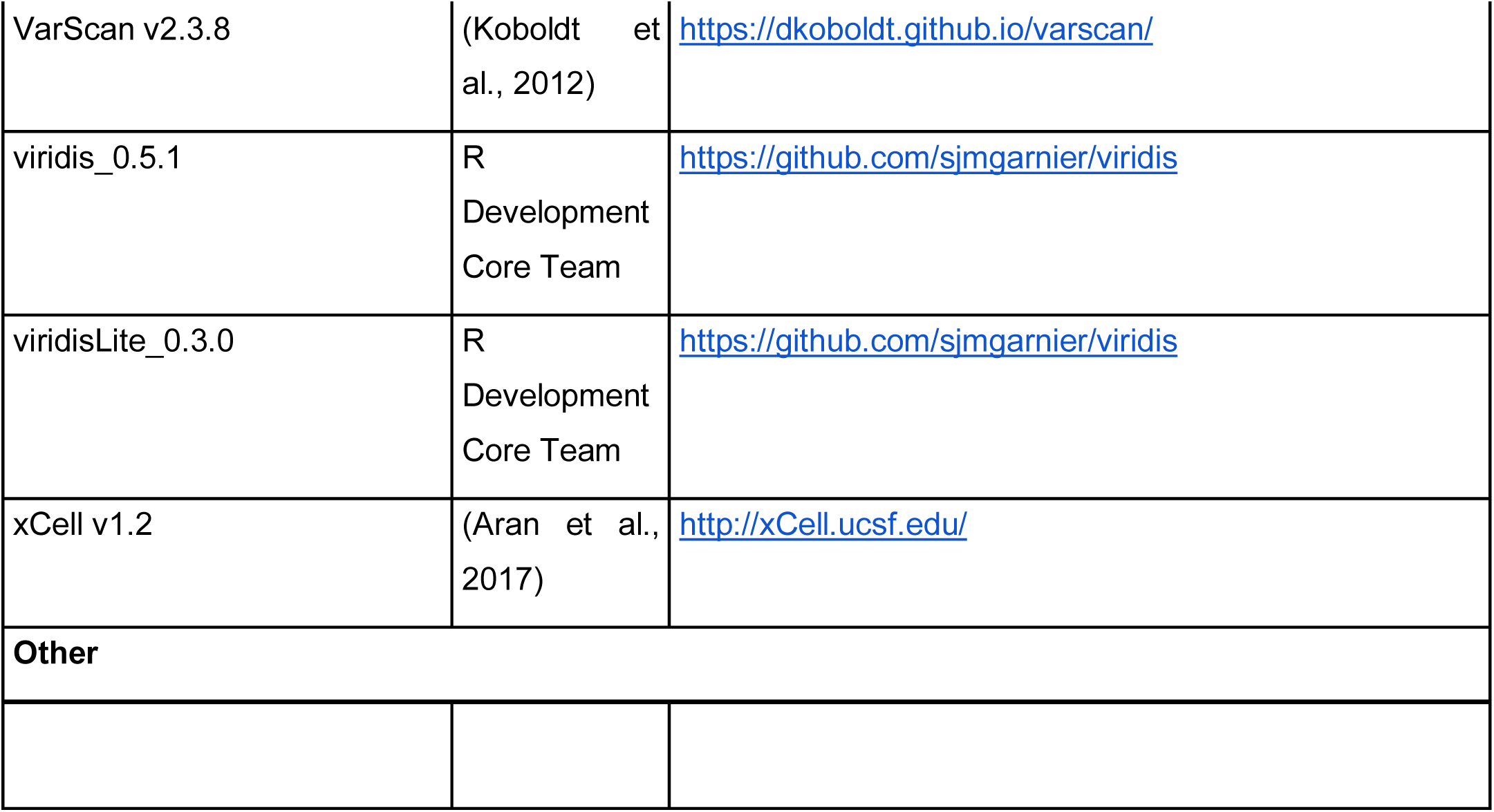

### EXPERIMENTAL MODEL AND SUBJECT DETAILS

#### Specimens and Clinical Data

All samples were collected with informed consent in concordance with Institutional Review Board (IRB) approval. Primary pancreatic adenocarcinoma samples were collected during surgical resection and verified by standard pathology (IRB protocol 201108117). Blood was collected at the time of surgery into vacuum tubes containing heparin or ethylenediaminetetraacetic acid (EDTA) (BD Bioscience). Cells were isolated by ficoll-density centrifugation and frozen in fetal bovine serum with 5% dimethyl sulfoxide.

Clinical data was captured in accordance with IRB protocol 20108117, at the time of informed consent and entered into the REDCap database.

#### Sample Processing

After verification by an attending pathologist, a 1.5 cm x 1.5 cm x 0.5 cm portion of the tumor was removed, photographed, weighed, and measured. Each piece was then subdivided into 6– 9 pieces (depending on the original size) and then further subdivided into four transverse cut pieces. Pieces were then placed into formalin, snap frozen in liquid nitrogen, DMEM, and formalin, respectively.

### METHOD DETAILS

#### Genomic DNA and RNA extraction

Tumor tissues and corresponding normal mucosae were obtained from surgically resected specimens, and after a piece was removed for fresh single-cell prep the remaining sample was snap-frozen in liquid nitrogen and stored at −80°C. Before bulk RNA/DNA extraction, samples were cryo-pulverized (Covaris) and aliquoted for bulk extraction methods. Genomic DNA was extracted from tissue samples with either the DNeasy Blood and Tissue Kit (Qiagen, 69504) or the QIAamp DNA Mini Kit (Qiagen, 51304). Total RNA was extracted with TRI reagent (Millipore Sigma, T9424) and treated with DNase I (Qiagen, 79254) using an RNeasy MinElute Cleanup Kit (Qiagen, 74204). RNA integrity was evaluated using either a Bioanalyzer (Agilent Technologies) or TapeStation (Agilent Technologies). Genomic germline DNA was purified from cryopreserved peripheral blood mononuclear cells (PBMCs) using the QiaAMP DNA Mini Kit (Qiagen, 51304) according to the manufacturer’s instructions (Qiagen, Valencia, CA). The DNA quantity was assessed by fluorometry using the Qubit dsDNA HS Assay (Q32854) according to manufacturer’s instructions (Thermo Fisher Scientific, Waltham, MA).

#### Whole-Exome Sequencing

100–250 ng of genomic DNA was fragmented on the Covaris LE220 instrument targeting 250bp inserts. Automated dual-indexed libraries were constructed with the KAPA Hyper library prep kit (Roche) on the SciClone NGS platform (Perkin Elmer). Up to ten libraries were pooled at an equimolar ratio by mass prior to the hybrid capture targeting a 5-µg library pool. The library pools were hybridized with the xGen Exome Research Panel v1.0 reagent (IDT Technologies) that spans a 39Mb target region (19,396 genes) of the human genome. The libraries were hybridized for 16–18 h at 65°C followed by stringent wash to remove spuriously hybridized library fragments. Enriched library fragments were eluted and PCR cycle optimization was performed to prevent over amplification. The enriched libraries were amplified with KAPA HiFi master mix (Roche) prior to sequencing. The concentration of each captured library pool was accurately determined through qPCR utilizing the KAPA library Quantification Kit according to the manufacturer’s protocol (Roche) to produce cluster counts appropriate for the Illumina NovaSeq-6000 instrument. 2x150 paired-end reads were generated targeting 12Gb of sequence to achieve ∼100x coverage per library.

#### RNA Sequencing

Total RNA integrity was determined using Agilent Bioanalyzer or 4200 Tapestation. Library preparation was performed with 500 ng to 1 ug of total RNA. Ribosomal RNA was blocked using FastSelect reagents (Qiagen) during cDNA synthesis. RNA was fragmented in reverse transcriptase buffer with FastSelect reagent and heated to 94°C for 5 min, 75°C for 2 min, 70°C for 2 min, 65°C for 2 min, 60°C for 2 min, 55°C for 2 min, 37°C for 5 min, 25°C for 5 min. mRNA was reverse transcribed to yield cDNA using SuperScript III RT enzyme (Life Technologies, per manufacturer’s instructions) and random hexamers. A second strand reaction was performed to yield ds-cDNA. cDNA was blunt ended, had an A base added to the 3’ ends, and then had Illumina sequencing adapters ligated to the ends. Ligated fragments were then amplified for 15 cycles using primers incorporating unique dual index tags. Fragments were sequenced on an Illumina NovaSeq-6000 S4 instrument generating approximately 30M paired end 2x150 reads per library.

#### Single-cell Suspension Preparation

For each tumor approximately 15–100 mg of 2–4 sections of each tumor and/or normal piece of tissue were cut into small pieces using a blade and processed separately. Enzymes and reagents from the human tumor dissociation kit (Miltenyi Biotec, 130-095-929) were added to the tumor tissue along with 1.75 mL of DMEM. The resulting suspension was loaded into a gentleMACS C-tube (Miltenyi Biotec, 130-093-237) and subject to the gentleMACS Octo Dissociator with Heaters (Miltenyi Biotec, 130-096-427). After 30–60 min on the heated dissociation program (37h_TDK_1), samples were removed from the dissociator and filtered through a 40-μm Mini-Strainer (PluriSelect #43-10040-60) or 40-μm Nylon mesh (Fisher Scientific, 22-363-547) into a 15-mL conical tube on ice. The sample was then spun down at 400 g for 5 min at 4 ℃. After removing the supernatant, when a red pellet was visible, the cell pellet was resuspended using 200 μL to 3 mL of ACK Lysis Solution (ThermoFisher, A1049201) for 1–5 min. To quench the reaction, 10 mL of PBS (Corning, 21-040-CM) with 0.5% BSA (Miltenyi Biotec, 130-091-376) was added and spun down at 400 g for 5 min at 4 ℃. After removing supernatant, the cells were resuspended in 1 mL of PBS with 0.5% BSA, and live and dead cells were visualized using Trypan Blue. If over 40% of dead cells were present, the sample was spun down at 400 g for 5 min at 4℃ and subject to the dead cell removal kit (Miltenyi Biotec, 130-090-101). Finally the sample was spun down at 400 g for 5 min at 4℃ and resuspended in 500 μL to 1 mL of PBS with 0.5% BSA to a final concentration of 700 to 1,500 cells per μL.

#### Single-cell library prep and sequencing

Utilizing the Chromium Next GEM Single Cell 3’ GEM, Library & Gel Bead Kit v3.1 and Chromium instrument, approximately 17,500 to 25,000 cells were partitioned into nanoliter droplets to achieve single-cell resolution for a maximum of 10,000 to 15,000 individual cells per sample (10x Genomics, 1000269). The resulting cDNA was tagged with a common 16nt cell barcode and 10nt Unique Molecular Identifier during the RT reaction. Full-length cDNA from poly-A mRNA transcripts was enzymatically fragmented and size-selected to optimize the cDNA amplicon size (approximately 400bp) for library construction (10x Genomics).The concentration of the 10x single-cell library was accurately determined through qPCR (Kapa Biosystems) to produce cluster counts appropriate for the HiSeq 4000 or NovaSeq 6000 platform (Illumina). 26x98bp sequence data were generated targeting 50K read pairs/cell, which provided digital gene expression profiles for each individual cell.

#### Proteomic and Phosphoproteomic Profiling Experiments

##### Protein Extraction and Lys-C/Trypsin Tandem Digestion

Tissue lysis and downstream sample preparation for global proteomic and phosphoproteomic analysis were carried out as previously described (Clark et al., 2019; Mertins et al., 2018). Approximately 25–50 mg of each cryo-pulverized HTAN tissue was resuspended in lysis buffer (8 M urea, 75 mM NaCl, 50 mM Tris, pH 8.0, 1 mM EDTA, 2 µg/mL aprotinin, 10 µg/mL leupeptin, 1 mM PMSF, 10 mM NaF, Phosphatase Inhibitor Cocktail 2 and Phosphatase Inhibitor Cocktail 3 [1:100 dilution], and 20 µM PUGNAc) by repeated vortexing. Lysates were clarified by centrifugation at 20,000 g for 10 min at 4°C, and protein concentrations were determined by BCA assay (Pierce). Proteins were reduced with 5 mM dithiothreitol (DTT, ThermoFisher) for 1 h at 37°C, and subsequently alkylated with 10 mM iodoacetamide (Sigma) for 45 min at room temperature (RT) in the dark. Samples were diluted 1:4 with 50 mM Tris-HCl (pH 8.0) and subjected to proteolytic digestion with LysC (Wako Chemicals) at 1 mAU:50 mg enzyme-to-substrate ratio for 2 h at RT, followed by the addition of sequencing grade modified trypsin (Promega) at 1:50 enzyme-to-substrate ratio and overnight incubation at RT. The digested samples were then acidified with 50% formic acid (FA, Fisher Chemicals) to pH 2. Tryptic peptides were desalted on reversed phase C18 SPE columns (Waters) and dried using Speed-Vac (Thermo Scientific).

##### TMT-11 Labeling of Peptides

Dried peptides from each sample were labeled with 11-plex TMT (Tandem Mass Tag) reagents (Thermo Fisher Scientific). 200 µg of peptides from each of the HTAN samples was dissolved in 80 µL of 100 mM HEPES, pH 8.5 solution. 30 HTAN samples were labeled in 3 TMT sets. A reference sample was created by pooling an aliquot from 26 HTAN samples (representing ∼90% of the sample cohort) and was included in all TMT 11-plex sets as a pooled reference channel (Channel 126). 5 mg of TMT reagent was dissolved in 500 µL of anhydrous acetonitrile, and then 30 µL of each TMT reagent was added to the corresponding aliquot of peptides. After 1 h incubation at RT, the reaction was quenched by incubation with 5% NH_2_OH for 15 min at RT. Following labeling, peptides were desalted on reversed phase C18 SPE columns (Waters) and dried using Speed-Vac (Thermo Scientific).

##### Peptide Fractionation by Basic Reversed-phase Liquid Chromatography (bRPLC)

To reduce the likelihood of peptides co-isolating and co-fragmenting due to high sample complexity, we employed extensive, high-resolution fractionation via basic reversed phase liquid chromatography (bRPLC). For each TMT set, about 2.2 mg of desalted peptides was reconstituted in 900 µL of 5 mM ammonium formate (pH 10) and 2% acetonitrile (ACN) and loaded onto a 4.6 mm x 250 mm RP Zorbax 300 A Extend-C18 column with 3.5-mm size beads (Agilent). Peptides were separated using an Agilent 1200 Series HPLC instrument using basic reversed-phase chromatography with Solvent A (2% ACN, 5 mM ammonium formate, pH 10) and a non-linear gradient of Solvent B (90% ACN, 5 mM ammonium formate, pH 10) at 1 mL/min as follows: 0% Solvent B (7 min), 0% to 16% Solvent B (6 min), 16% to 40% Solvent B 60 min), 40% to 44% Solvent B (4 min), 44% to 60% Solvent B (5 min) and then held at 60% Solvent B (14 min). Collected fractions were concatenated into 24 fractions as described previously (Mertins et al., 2018); 5% of each of the 24 fractions was aliquoted for global proteomic analysis, dried down in a Speed-Vac, and resuspended in 3% ACN, 0.1% formic acid prior to ESI-LC-MS/MS analysis. The remaining sample was utilized for phosphopeptide enrichment.

##### Enrichment of Phosphopeptides by Fe-IMAC

The remaining 95% of the fractions were further concatenated into 12 fractions prior to phosphopeptide enrichment using immobilized metal affinity chromatography (IMAC) as previously described (Mertins et al., 2018). In brief, Ni-NTA agarose beads were utilized to prepare Fe^3+^-NTA agarose beads, and then about 200 µg of peptides of each fraction reconstituted in 80% ACN/0.1% trifluoroacetic acid were incubated with 10 µL of the Fe^3+^-IMAC beads for 30 mins. Samples were then spun down, and the supernatant containing unbound peptides was removed. The beads were brought up in 80% ACN, 0.1% trifluoroacetic acid and then loaded onto equilibrated C-18 Stage Tips, and washed by 80% ACN, 0.1% trifluoroacetic acid, rinsed twice with 1% formic acid, followed by sample elution off the Fe^3+^-IMAC beads with 100 µL of 500 mM dibasic potassium phosphate, pH 7.0. C-18 Stage Tips were then washed twice with 1% formic acid, followed by elution of the phosphopeptides from the C-18 Stage Tips with 80 µl of 50% ACN, 0.1% formic acid twice. Samples were dried down and resuspended in 3% ACN, 0.1% formic acid prior to ESI-LC-MS/MS analysis.

##### ESI-LC-MS/MS for Global Proteome and Phosphoproteome Analysis

The global proteome and phosphoproteome fractions were analyzed as described in a previous study (Clark et al., 2019). Peptides (∼0.8 µg) were separated on an Easy nLC 1200 UHPLC system (Thermo Scientific) on an in-house packed 20 cm x 75 mm diameter C18 column (1.9 mm Reprosil-Pur C18-AQ beads (Dr. Maisch GmbH); Picofrit 10 mm opening (New Objective)). The column was heated to 50°C using a column heater (Phoenix-ST). The flow rate was 0.300 μL/min with 0.1% formic acid and 2% acetonitrile in water (A) and 0.1% formic acid, 90% acetonitrile (B). The global peptides were separated with a 6–30% B gradient in 84 mins and analyzed using the QE-HFX (Thermo Scientific). Parameters were as follows MS1: resolution – 120,000, mass range – 400 to 2000 m/z, RF Lens – 30%, AGC Target 3e6, Max IT – 50 ms, charge state include -2-5, dynamic exclusion – 20 s, top 20 ions selected for MS2; MS2: resolution – 45,000, collision energy NCE – 32, isolation width (m/z) – 0.7, AGC Target – 1.0e5, Max IT – 96 ms. The phosphopeptides were separated with a 6–30% B gradient in 84 mins and analyzed using the Lumos (Thermo Scientific). Parameters were as follows MS1: resolution – 60,000, mass range – 350 to 1800 m/z, RF Lens – 30%, AGC Target 4.0e5, Max IT – 50 ms, charge state include -2-6, dynamic exclusion – 45 s, top 20 ions selected for MS2; MS2: resolution – 50,000, HCD collision energy – 34, isolation width (m/z) – 0.7, AGC Target – 2.0e5, Max IT – 100 ms.

### QUANTIFICATION AND STATISTICAL ANALYSIS

#### Genomic Data Analysis

##### Somatic Variant Calling

Somatic variants were called from whole-exome tumor-normal paired BAMs using somaticwrapper v1.5, a pipeline designed for detection of somatic variants from tumor and normal whole-exome sequence (WES) data. The pipeline merges and filters variant calls from four callers: Strelka v2.9.2 (Kim et al., 2018), VarScan v2.3.8 (Koboldt et al., 2012), Pindel v0.2.5 (Ye et al., 2009), and MuTect v1.1.7 (Cibulskis et al., 2013). SNV calls were obtained from Strelka, Varscan, and MuTect. Indel calls were obtained from Strelka, Varscan, and Pindel. The following filters were applied to obtain variant calls of high confidence: normal VAF ≤ 0.02 and tumor VAF ≥ 0.05, read depth in tumor ≥ 14 and normal ≥ 8, indel length < 100 bp, all variants must be called by 2 or more callers, all variants must be exonic, and variants in dbSNP but not in COSMIC excluded.

##### KRAS Hotspot Genotyping

To verify manually and/or determine the KRAS mutation status at KRAS hotspots G12, G13, and Q61, we used bam-readcount. For each case, we first applied bam-readcount to generate readcounts for each of the 9 bases in these loci and then calculated VAF values of all the KRAS hotspots based on reference and alternative base read counts at each position.

##### Germline Variant Calling and Annotation

Germline variant calling was performed using an in-house pipeline, germlinewrapper v1.1, which implements multiple tools for the detection of germline INDELs and SNVs. Germline SNVs were identified using VarScan v2.3.8 (with parameters: --min-var-freq 0.10 --p-value 0.10 --min-coverage 3 --strand-filter 1) operating on a mpileup stream produced by samtools v1.2 (with parameters: -q 1 -Q 13) and GATK v4.0.0.0 (McKenna et al., 2010) using its haplotype caller in single-sample mode with duplicate and unmapped reads removed and retaining calls with a minimum quality threshold of 10. All resulting variants were limited to the coding regions of the full-length transcripts obtained from Ensembl release 95 plus an additional two base pairs flanking each exon to cover splice donor/acceptor sites. We required variants to have allelic depth ≥ 5 reads and alternative allele frequencies ≥ 20% in both the tumor and normal samples. We used bam-readcount v0.8 for reference and alternative alleles quantification (with parameters: -q 10 -b 15) in both normal and tumor samples. Additionally, we filtered all variants with ≥ 0.05% frequency in gnomAD v2.1 (Karczewski et al., 2017) and The 1000 Genomes Project (The 1000 Genomes Project Consortium, 2015).

##### Germline Variant Pathogenic Classification

For annotation and prioritization of the filtered germline variants, we used our automatic variant classification tool CharGer v0.5.4 (Scott et al., 2019), which computes a classification score based on ACMG-AMP guidelines. CharGer automatically marks as pathogenic those input variants that are marked as known pathogenic in ClinVar’s curated database and marks as likely pathogenic those variants with a CharGer score > 8. All pathogenic or likely pathogenic variants had both their normal and tumor samples reviewed manually by us using the Integrative Genomics Viewer (IGV) software.

#### scRNA-Seq Quantification and Analysis

##### scRNA-Seq Data Preprocessing

For each sample, we obtained the unfiltered feature-barcode matrix per sample by passing the demultiplexed FASTQs to Cell Ranger v3.1.0 ‘count’ command using default parameters and the prebuilt GRCh38 genome reference v3.0.0 (GRCh38 and Ensembl 93). Seurat v3.1.2 (Butler et al., 2018; Hafemeister and Satija, 2019) was used for all subsequent analysis. First, a series of quality filters was applied to the data to remove those barcodes which fell into any one of these categories recommended by Seurat: too few total transcript counts (< 300); possible debris with too few genes expressed (< 200) and too few UMIs (< 1,000); possible more than one cell with too many genes expressed (> 10,000) and too many UMIs (> 10,000); possible dead cell or a sign of cellular stress and apoptosis with too high proportion of mitochondrial gene expression over the total transcript counts (> 10%). We constructed a Seurat object using the unfiltered feature-barcode matrix for each sample. Each sample was scaled and normalized using Seurat’s ‘SCTransform’ function to correct for batch effects (with parameters: vars.to.regress = c(“nCount_RNA”, “percent.mito”), variable.features n = 2000). Any merged analysis or subsequent subsetting of cells/samples underwent the same scaling and normalization method. Cells were clustered using the original Louvain algorithm (Blondel et al., 2008) and top 20 PCA dimensions via ‘FindNeighbors’ and ‘FindClusters’ (with parameters: resolution = 0.5) functions. The resulting merged and normalized matrix was used for the subsequent analysis.

##### scRNA-Seq Cell Type Annotation

Main cell types were assigned to each cluster by manually reviewing the expression of a comprehensive set of marker genes. These assignments were all done by one person to maximize consistency. The marker genes used were *KRT19, KRT8, KRT18, KRT17, KRT7, KRT5, KRT6A, KRT14, EPCAM, TACSTD2, ANXA2, S100A10, S100A11, S100A16, TPM1, TFF1, S100A6, AGR2, C19orf33* (tumor); *INS, GCG, SST, GHR, PPY, GCK, PCSK1, PCSK2, CHGA, CHGB, SYP, KCNJ11* (islet); *CTRB1, CELA3A, CELA3B, CTRB2, PLA2G1B, PRSS2, SPINK1, CLPS, CPA1, PRSS1, CPA2, REG1A, PNLIP, SYCN, PNLIPRP1, CTRC, KLK1, CELA2A, CPB1* (acinar); *VWF, PECAM1, FLT4, FLT1, FLT3, KDR, PLVAP, ANGPT2, TRIM24, ACTA2* (endothelial); *TIMP1, FN1, POSTN, ACTA2, BST2, LY6D, COL6A1, SLC20A1, COL6A2, KRT16, CD9, S100A4, EMP1, LRRC8A, EPCAM, PDPN, ITGB1, PDGFRA, THY1* (fibroblast); *HBD, GYPA, HBA1, HBA2, CA1, HBB, BRSK1* (erythrocyte); *SDC1, IGHG1, IGHG3, IGHG4* (plasma); *CD19, MS4A1, CD79A, CD79B, CD83, CD86* (B cells); *CD8B, CD8A, CD3E, CD3D* (CD8 T cells); *CD8B, CD8A, CD4* (CD4 T cells); *XCL2, XCL1, SPON2, KLRF1, KIR2DL3, IL2RB, HOPX, CLIC3, CD7, KLRB1, KLRD1, GZMA, PRF1, CD160, NCAM1, FCGR3A* (NK); *FCER1A, KIT, FCER2, ENPP3* (mast); *CD1C, CD1A, FLT3, ZBTB46, XCR1, CLEC9A, IRF8, FLT3, ZBTB46, BATF3* (cDCs); *LY6G6D, MPO, FUT4, FCGR3A* (neutrophils); *ITGAM, LGALS3, CD68, CD163, LYZ, ADGRE1, LAMP2* (macrophages); *CD14, FCGR3A, FCGR1A* (monocytes). We further subdivided certain cell types into subtypes or cell states using the following: *IKZF2, TNFRSF18, FOXP3, CTLA4, IL7R, IL2RA* (Treg); *GZMH, GZMK, GZMB, GZMA, PRF1, IFNG, FASLG, LAMP1, CD8A, CD8B, CD3E, CD3D* (cytotoxic T cell); *VSIR, TIGIT, ICOS, EOMES, HAVCR2, PDCD1, BTLA, CD244, LAG3, CD160, CTLA4, CD96* (exhausted T cell); *MKI67* (proliferation marker); *ACTA2, FAP, PDPN, PDGFRA* (general CAF); *IL6, CXCL1, CXCL12, CXCL2* (iCAF); *ACTA2, THY1, TAGLN* (myCAF); *CD74, SAA2, SAA1* (apCAF). For ADM populations, we used a combination of tumor and acinar markers.

##### scRNA-Seq Subtype Assignment

We used 3 bulk gene marker sets (Bailey, Moffitt, and Collisson) and their respective subtypes for scRNA-Seq PDAC cluster assignment. For each subtype from the Bailey and Moffitt gene sets, the top 20 genes were selected, while from the Collisson gene set, all genes were used due to the lower marker count per subtype. The resulting gene marker lists grouped by subtype are as follows. Bailey set: *MTMR7, ARX, ABCC8, CEACAM7, CACNA2D2, MAPK8IP2, SYT7, LTF, NR1H4, SLC7A14, PRSS3, CPA1, AQP8, FGL1, SERPINI2, REG1A, NR5A2, RBPJL, KIRREL2, CLPS* (Adex); *CFH, CD99, HECW1, PLXND1, RECQL, C1orf112, M6PR, FKBP4, GGCT, DBF4, SEMA3F, ANKIB1, MAD1L1, CDC27, WDR54, DPM1, NME2, UTP18, SLC25A39, TTC27* (Squamous); *ENPP4, CFTR, CYP51A1, ABP1, LASP1, TMEM176A, ICA1, DBNDD1, CASP10, SARM1, UPF1, ACSM3, SPPL2B, PDK2, OSBPL7, TMEM98, BAIAP2L1, ALDH3B1, TTC22, FARP2* (PancProg); *CD38, PDK4, FBXL3, CD79B, MYLIP, RWDD2A, ACPP, TRAF3IP3, ACAP1, ARHGAP15, CST7, P2RY10, SIRPG, GRAP2, FGR, ITGAL, CEACAM21, CD4, BTK, TYROBP* (Immunogenic). Collisson set: *REG1B, REG3A, REG1A, PNLIPRP2, CEL, PNLIP, PLA2G1B, CELA3A, CPB1, CELA3B, CTRB2, CLPS, CELA2B, PRSS2, PRSS1, GP2, SLC3A1, CFTR, SLC4A4, SPINK1* (Exocrine-like); *AIM2, CALHM6, GPM6B, S100A2, KRT14, CAV1, LOX, SLC2A3, TWIST1, PAPPA, NT5E, CKS2, HMMR, SLC5A3, PMAIP1, PHLDA1, SLC16A1* (Quasi-Mesenchymal); *FERMT1, HK2, AHNAK2, TMEM45B, SDR16C5, GPRC5A, AGR2, S100P, FXYD3, ST6GALNAC1, CEACAM5, CEACAM6, TFF1, TFF3, CAPN8, FOXQ1, ELF3, ERBB3, TSPAN8, TOX3, LGALS4, PLS1, GPX2, ATP10B, MUC13* (Classical). Moffitt set: *S100A2, KRT6A, CST6, GPR87, SCEL, ANXA8L2, KRT6C, SERPINB4, SERPINB3, LY6D, LY6D, PLAG1, IL20RB, C16orf74, DCBLD2, KRT17P3, HMGA2, SPRR3, SPRR1B, KRT17* (Basal-like); *TM4SF5, TFF1, PLA2G10, GPX2, SPINK4, LOC400573, BTNL8, DMBT1, ATAD4, TFF3, FAM3D, LYZ, MYO1A, ANXA10, CLRN3, AKR1B10, CTSE, TSPAN8, LGALS4, REG4* (Classical). To assign the scRNA-Seq subtype, we calculated the “subtype score” for each cluster identified in the merged Seurat object described in the “scRNA-Seq Data Preprocessing” section. The subtype scores were defined by taking an average of selected bulk markers for each subtype in each marker set. Each cluster was then assigned to the subtype with the highest score within each marker set.

##### Composite Heterogeneity Score

For the single cell-based composite heterogeneity score, we first calculated the mean pairwise Spearman correlation of cell type proportions between samples from each case for a case-level cell type proportion statistic. We then calculated the average expression of all genes for each sample and obtained the mean pairwise Spearman correlation across all genes between samples from each case to obtain a case-level expression statistic. We defined the mean of these two scores, subtracted from 1, as the scRNA-based composite heterogeneity score for each patient. For the heterogeneity scores at the cell-type level, the score is calculated by correlating average expression between cells from each cell type only.

##### scVarScan Mutation Mapping

We applied our in-house tool scVarScan that can identify reads supporting the reference and variant alleles covering the variant site in each individual cell by tracing cell and molecular barcode information in an scRNA bam file. For mapping, we used high-confidence somatic mutations from WES data. Additionally, we use cancerhotspots.org (Chang et al., 2018) to obtain the most common KRAS hotspot mutations at G12, G13, and Q61 and use scVarScan to detect potential minority KRAS mutations in each sample.

##### Single-cell RNA CNV Detection

To detect large-scale chromosomal copy number variations using single-cell RNA-seq data, inferCNV (version 0.8.2) was used with default parameters recommended for 10x Genomics data. All cells that are not tumor were pooled together for the reference normal set. InferCNV was run at a sample level and only with post-QC filtered data. In order to calculate arm-level CNV events, we used an in-house script to match the gene-level inferCNV output to chromosome bands and take the mean value for each arm.

##### Differential scRNA Expression Analyses

For cell-level and cluster-level differential expression, we used the ’FindMarkers’ or “FindAllMarkers’ Seurat function as appropriate, using a minimum percent of 0.25 (parameter min.pct = 0.25) and looking only in the positive direction, as lack of expression is harder to interpret due to the sparsity of the data. The resulting DEGs were then filtered for adjusted p-value < 0.05 and sorted by fold change. All differential expression analyses were carried out using the “SCT” assay.

##### Acinar and Ductal Scores

To create gene module scores, we used the Seurat function “AddModuleScore” with default parameters and input our acinar or ductal marker genes lists as follows: *KRT19, KRT8, KRT18, KRT17, KRT7, KRT5, KRT6A, KRT14, EPCAM, TACSTD2, ANXA2, S100A10, S100A11, S100A16, TPM1, TFF1, S100A6, AGR2, C19orf33* (tumor); and *CTRB1, CELA3A, CELA3B, CTRB2, PLA2G1B, PRSS2, SPINK1, CLPS, CPA1, PRSS1, CPA2, REG1A, PNLIP, SYCN, PNLIPRP1, CTRC, KLK1, CELA2A, CPB1* (acinar).

##### Receptor Ligand Interactions

We used the CellPhoneDB tool (Efremova et al., 2020) in order to detect significant pairs of receptor-ligand interactions between cell types. This comparison was done at the sample level using default parameters between tumor and lymphocyte cell types.

##### Cell Surface Annotation

To annotate a given biomarker, we annotated each DEG by its subcellular location. Three databases were used to curate the subcellular location information: 1) Gene Ontology Term 0005886; 2) Mass Spectrometric-Derived Cell Surface Protein Atlas (CSPA) (Bausch-Fluck et al., 2015); 3) The Human Protein Atlas (HPA) subcellular location data based on HPA version 19.3 and Ensembl version 92.38.

#### Bulk Proteogenomic Analyses

##### DNA and RNA Sample QC

Bulk sequencing data quality metrics (adaptor content, mapping quality, coverage, and contamination/swaps/mislabeling) were determined for DNA and RNA bams using our in-house pipeline SeqQEst. The inclusion criteria for paired DNA and RNA bams was that there be no contamination and sufficient coverage (> 50x coding region coverage in WES or > 50Mb mapped depth in RNA-Seq data).

##### RNA Quantification

We used our in-house bulk-RNA-Seq expression analysis pipeline for quantification. Briefly, for each sample, the raw sequence reads were aligned into BAM files using STAR (Dobin et al., 2013) (version 2.7.4a) 2-pass alignment with GRCh38 as the reference. The resulting BAM files were then quantified as a raw count-matrix using read feature counts using Subread (Liao et al., 2013) (version 2.0.1). For both alignment and quantification, gene annotations were based on Gencode v34. The raw counts were then converted to FPKM-UQ based on GDC’s formula (https://docs.gdc.cancer.gov/Data/Bioinformatics_Pipelines/Expression_mRNA_Pipeline/#upper-quartile-fpkm) and then log2 transformed with 1 pseudocount.

##### Proteomic and Phosphoproteomics Quantification

Proteomic data processing followed the methods detailed in Clark et al., 2019. Briefly, raw mass spectrometry files were converted into open mzML format, then searched using the MSFragger database against a RefSeq protein sequence database appended with an equal number of decoy sequences. The specific parameters and software are detailed in the Clark et al., 2019 study.

##### Heterogeneity Score

Similar to the scRNA average expression metric, for each of the bulk RNA, proteomic, and phosphoproteomic data, we calculated the mean pairwise Spearman correlation across all genes between samples from each case to obtain a case-level correlation metric. We then defined the heterogeneity score as this mean correlation subtracted from 1. For phosphoproteomics, this was done using phosphosite abundances, not phosphoprotein abundances.

##### Expression-Based Subtyping

Bulk expression subtyping is done by selecting top genes from each gene marker set (Bailey, Moffitt, and Collisson), normalizing expression, and clustering with the ConsensusClusterPlus (Wilkerson and Hayes, 2010) package in R. The top gene selection process is the same as that described in the scRNA-Seq subtype assignment method section. The log2 upper quartile normalized FPKM reads of these genes were standardized with z-score scaling, making each gene’s mean and standard deviation equal to 0 and 1, respectively, thereby allowing all genes on different expression levels to contribute equally to clustering. These normalized expression matrices were then clustered using ConsensusClusterPlus (with parameters for 1000 iterations and maximum k = 8). The optimum k for each marker set was selected based on the number of subtypes originally identified in each gene marker set (2, 3, and 4 for Moffitt, Collisson, and Bailey, respectively.) The clusters were then assigned to each subtype by inspecting the expression level of their respective marker gene lists.

##### Mutation Impact on the Proteome and Phosphoproteome

We used an aggregated database of interacting proteins that combines Omnipath, DEPOD, CORUM, Signor2, and Reactome databases as previously described (Dou et al., 2020). We focused our analyses on PDAC SMGs previously reported in the literature, but only KRAS and TP53 had large enough numbers in each comparison group for sufficient statistical power (Bailey et al., 2018). For each interacting pair, we split samples with and without mutations in partner A and compared expression levels (both protein and phosphosites) both in *cis* (partner A) and in *trans* (partner B). We calculated the median difference in expression and tested for significance using Wilcoxon rank sum tests, with Benjamini-Hochberg multiple test correction. We further refined the list of *trans* interactions by filtering proteins that are not part of oncogenic processes identified in TCGA (Sanchez-Vega et al., 2018).

##### KRAS Phosphosignaling Analysis

Oncogenic KRAS signaling in PDAC is believed to pass through three major pathways: Raf/Mek/Erk, PI3K/Pdk1/Akt, and the Ral guanine nucleotide exchange factor pathway (Eser et al., 2014). We focused on the Raf/Mek/Erk pathway (along with PI3K/Pdk1/Akt) because its signaling is controlled by phosphorylation. We used the phosphosites identified in this pathway as detailed in Collisson et al. (2012).

##### Differential Proteogenomic Analysis

For differential analysis between groups of samples using bulk data (gene expression, proteomics, and phosphoproteomics), we used Wilcoxon rank-sum tests to test for differential abundances of genes, proteins, and phosphosites. We required that at least 50% of samples in each comparison group have non-missing values. P-values were then adjusted using the Benjamini-Hochberg multiple test correction to obtain features with an FDR cutoff ≥ 0.05.

##### Immune Clustering Using Cell Type Enrichment Scores

The abundance of each cell type was inferred by the xCell web tool (Aran et al., 2017), which performed the cell type enrichment analysis from gene expression data for 64 immune and stromal cell types (default xCell signature). xCell is a gene signatures-based method learned from thousands of pure cell types from various sources. We used the FPKM-UQ expression matrix as the input to xCell. xCell generated an immune score per sample that integrates the enrichment scores of various cell types, including B cells, CD4+ T-cells, CD8+ T-cells, DC, eosinophils, macrophages, monocytes, mast cells, neutrophils, and NK cells. Immune subtypes of HTAN PDAC cohorts were generated based on the consensus clustering of the xCell cell type enrichment scores (Wilkerson and Hayes, 2010). Among the 64 cell types tested in xCell, we selected cell types with at least 2 samples with xCell enrichment p < 0.01 and performed the consensus immune clustering based on the z-score normalized xCell enrichment scores. The consensus clustering was determined by the R package ConsensusClusterPlus (parameters: reps = 2000, pItem = 0.9, pFeature = 0.9, clusterAlg = “kmdist”, distance = “spearman”).

##### ESTIMATE Immune and Stroma Scores

The ESTIMATE scores reflecting the overall immune and stromal infiltration and tumor purity estimation were calculated by the R package ESTIMATE (Yoshihara et al., 2013) using the normalized RNA expression data (FPKM-UQ). The ESTIMATE algorithm is based on single-sample gene set enrichment analysis and generates three scores: 1) stromal score (which captures the presence of stroma in tumor tissue), 2) immune score (which represents the infiltration of immune cells in tumor tissue), and 3) estimate score (which infers tumor purity).

##### CiVIC Drug Matching

We obtained evidence of expression-based response to drugs from CIViC (Griffith et al., 2017). We filtered the database for only sensitive results and positive direction (i.e., “expression” and “overexpression”). We then matched these annotations to upregulated DEGs in our comparison groups. In the present study we used the 06/02/2020 nightly clinical evidence summary annotations.

#### Imaging Mass Cytometry

##### Tissue Staining for IMC

Two 5-μm thick sections from each pancreas piece were stained with the full antibody panel (see Key Resources Table). Slides were placed into a 60°C oven for 2 h, deparaffinized with xylene, and then rehydrated with a series of alcohols of decreasing concentrations. Antigen retrieval was performed by placing slides into a pre-warmed solution of Tris-EDTA, pH 9.0 and incubated at 96°C for 30 min. Slides were then cooled and washed in water and PBS. Non-specific staining was blocked in PBS with 3% Bovine Serum Albumin (BSA). A cocktail containing all antibodies was prepared in PBS with 3% BSA, and slides were stained overnight at 4°C. Slides were then washed with PBS/0.2% Triton-X and incubated with an iridium intercalator for 30 min at room temperature. Ruthenium red counterstain was applied for 10 min after which slides were washed in diH2O then air-dried for 20 min.

##### IMC Acquisition

Regions of interest (ROI) were selected based on hematoxylin and eosin (H&E) stains on adjacent sections. Selected areas of 1,000 μ x 1,000 μ were ablated by a laser which rasters across the selected ROI at a rate of 200 Hz (200 pixels/s). The time needed for complete ablation of each ROI selected is about 2 h. The ablated tissue is then carried by an inert stream of helium and argon gas into the Helios (Fluidigm, South San Francisco CA) where the material is completely ionized in inductively coupled plasma. The ionized material enters into a time of flight (TOF) detector where it is separated based on mass. Laser shot signals are recorded in order and generate a pseudo-colored intensity map of each mass channel. These data are then exported as MCD files.

##### IMC Analyses

MCD images were converted to 32-bit ome.tiff images with Fluidigm MCD Viewer. Nuclear segmentation was performed with Halo(tm) multiplex imaging analysis software. Cell boundaries were delineated by expanding the nuclear segmentation boundary for each cell by 3 pixels. Marker intensities for each cell were then exported from Halo and normalized with the following steps: 1) division by cellular area, 2) capping to the 99% percentile, and 3) scaling by standard deviation and mean-centering. Marker intensities were then decomposed with PCA and UMAP, and nearest neighbors (n=15) were identified for each cell. Cells were then clustered with the Leiden algorithm (implemented by the Python library Scanpy). Tumor cell clusters were identified as those harboring elevated levels of Pan-Keratin. Cells from this tumor cluster were then annotated according to their Pan-Keratin and Alpha-Amylase intensities. The cells were split into four partitions, (Alpha-Amylase+, Pan-Keratin+), (Alpha-Amylase+, Pan-Keratin-), (Alpha-Amylase-, Pan-Keratin+), and (Alpha-Amylase-, Pan-Keratin-) based on the following intensity thresholds: Alpha-Amylase < 2 or ≥ 2, and Pan-Keratin < 2 or ≥ 2.

#### Additional Methods

##### Pathological Parameters and Assessment

Each tumor that is subdivided into smaller increments is subjected to H&E staining and assessed by a pathologist for the following parameters: percentage of viable tumor present, tumor differentiation, presence of recognizable pancreatic parenchyma surrounding or interspersed between tumor, lymphovascular invasion, and perineural invasion. Both slices of each tumor piece, both L1 and L4 when available, were subjected to assessment. For the correlation with scRNA-based tumor percentages, we averaged the top and bottom slide (L1 and L4) tumor estimates.

##### Pathway analysis

For each comparison, we obtained the top 30 genes ranked by highest fold change that are significantly different between the comparison groups (FDR < 0.05). We used ConsensusPathDB-human for gene set over-representation analysis (Kamburov et al., 2013).

### DATA AND CODE AVAILABILITY

All raw image and sequencing data will be deposited into the publicly available HTAN DCC at https://humantumoratlas.org/htan-dcc/.

## ADDITIONAL RESOURCES SUPPLEMENTAL TABLE LEGENDS

**Table S1**: Cohort information including key clinical and molecular phenotypes.

**Table S2**: Bulk omics data including somatic and germline variants and proteogenomics data.

**Table S3:** Tumor biomarkers and tumor DEGs between treatment groups.

## REFERENCES

1000 Genomes Project Consortium. (2015). A global reference for human genetic variation. Nature, 526(7571), 68–74.

Alizadeh, A. A., Aranda, V., Bardelli, A., Blanpain, C., Bock, C., Borowski, C., … & Esteller, M. (2015). Toward understanding and exploiting tumor heterogeneity. Nature medicine, 21(8), 846.

American Cancer Society (2020). Cancer Facts & Figures 2020. Atlanta: American Cancer Society.

Aran, D., Hu, Z., & Butte, A. J. (2017). xCell: digitally portraying the tissue cellular heterogeneity landscape. Genome biology, 18(1), 1–14.

Bailey, M. H., Tokheim, C., Porta-Pardo, E., Sengupta, S., Bertrand, D., Weerasinghe, A., … & Ng, P. K. S. (2018). Comprehensive characterization of cancer driver genes and mutations. Cell, 173(2), 371–385.

Bailey, P., Chang, D. K., Nones, K., Johns, A. L., Patch, A. M., Gingras, M. C., … & Nourse, C. (2016). Genomic analyses identify molecular subtypes of pancreatic cancer. Nature, 531(7592), 47–52.

Balachandran, V. P., Beatty, G. L., & Dougan, S. K. (2019). Broadening the impact of immunotherapy to pancreatic cancer: challenges and opportunities. Gastroenterology, 156(7), 2056–2072.

Bausch-Fluck, D., Hofmann, A., Bock, T., Frei, A. P., Cerciello, F., Jacobs, A., … & Schiess, R. (2015). A mass spectrometric-derived cell surface protein atlas. PloS one, 10(4), e0121314.

Biffi, G., Oni, T. E., Spielman, B., Hao, Y., Elyada, E., Park, Y., … & Tuveson, D. A. (2019). IL1-induced JAK/STAT signaling is antagonized by TGFβ to shape CAF heterogeneity in pancreatic ductal adenocarcinoma. Cancer discovery, 9(2), 282–301.

Birnbaum, D. J., Finetti, P., Lopresti, A., Gilabert, M., Poizat, F., Turrini, O., … & Mamessier, E. (2016). Prognostic value of PDL1 expression in pancreatic cancer. Oncotarget, 7(44), 71198.

Blondel, V. D., Guillaume, J. L., Lambiotte, R., & Lefebvre, E. (2008). Fast unfolding of communities in large networks. Journal of statistical mechanics: theory and experiment, 2008(10), P10008.

Butler, A., Hoffman, P., Smibert, P., Papalexi, E., & Satija, R. (2018). Integrating single-cell transcriptomic data across different conditions, technologies, and species. Nature biotechnology, 36(5), 411–420.

Chang, M. T., Bhattarai, T. S., Schram, A. M., Bielski, C. M., Donoghue, M. T., Jonsson, P., … & Gorelick, A. (2018). Accelerating discovery of functional mutant alleles in cancer. Cancer discovery, 8(2), 174–183.

Chatterjee, M., Ben-Josef, E., Thomas, D. G., Morgan, M. A., Zalupski, M. M., Khan, G., … & Bekaii-Saab, T. (2015). Caveolin-1 is associated with tumor progression and confers a multi-modality resistance phenotype in pancreatic cancer. Scientific reports, 5, 10867.

Chen, D., & Che, G. (2014). Value of caveolin-1 in cancer progression and prognosis: Emphasis on cancer-associated fibroblasts, human cancer cells and mechanism of caveolin-1 expression. Oncology letters, 8(4), 1409–1421.

Chung, W., Eum, H. H., Lee, H. O., Lee, K. M., Lee, H. B., Kim, K. T., … & Kan, Z. (2017). Single-cell RNA-seq enables comprehensive tumour and immune cell profiling in primary breast cancer. Nature communications, 8(1), 1–12.

Clark, D. J., Dhanasekaran, S. M., Petralia, F., Pan, J., Song, X., Hu, Y., … & Ma, W. (2019). Integrated proteogenomic characterization of clear cell renal cell carcinoma. Cell, 179(4), 964–983.

Collisson, E. A., Sadanandam, A., Olson, P., Gibb, W. J., Truitt, M., Gu, S., … & Feiler, H. S. (2011). Subtypes of pancreatic ductal adenocarcinoma and their differing responses to therapy. Nature medicine, 17(4), 500–503.

Collisson, E. A., Trejo, C. L., Silva, J. M., Gu, S., Korkola, J. E., Heiser, L. M., … & Spellman, P. T. (2012). A central role for RAF→MEK→ERK signaling in the genesis of pancreatic ductal adenocarcinoma. Cancer discovery, 2(8), 685–693.

Conroy, T., Hammel, P., Hebbar, M., Ben Abdelghani, M., Wei, A. C., Raoul, J. L., … & Lecomte, T. (2018). FOLFIRINOX or gemcitabine as adjuvant therapy for pancreatic cancer. New England Journal of Medicine, 379(25), 2395–2406.

DiMagno, E. P., Malagelada, J. R., & Go, V. L. (1979, March). The relationships between pancreatic ductal obstruction and pancreatic secretion in man. In Mayo Clinic Proceedings (Vol. 54, No. 3, p. 157).

Dou, Y., Kawaler, E. A., Zhou, D. C., Gritsenko, M. A., Huang, C., Blumenberg, L., … & Liu, W. (2020). Proteogenomic characterization of endometrial carcinoma. Cell, 180(4), 729–748.

Eckert, F., Schilbach, K., Klumpp, L., Bardoscia, L., Sezgin, E. C., Schwab, M., … & Huber, S. M. (2018). Potential role of CXCR4 targeting in the context of radiotherapy and immunotherapy of cancer. Frontiers in Immunology, 9, 3018.

Efremova, M., Vento-Tormo, M., Teichmann, S. A., & Vento-Tormo, R. (2020). CellPhoneDB: inferring cell–cell communication from combined expression of multi-subunit ligand–receptor complexes. Nature protocols, 15(4), 1484–1506.

Elyada, E., Bolisetty, M., Laise, P., Flynn, W. F., Courtois, E. T., Burkhart, R. A., … & Sivajothi, S. (2019). Cross-species single-cell analysis of pancreatic ductal adenocarcinoma reveals antigen-presenting cancer-associated fibroblasts. Cancer discovery, 9(8), 1102–1123.

Eser, S., Schnieke, A., Schneider, G., & Saur, D. (2014). Oncogenic KRAS signalling in pancreatic cancer. British journal of cancer, 111(5), 817–822.

Feng, M., Xiong, G., Cao, Z., Yang, G., Zheng, S., Song, X., … & Zhao, Y. (2017). PD-1/PD-L1 and immunotherapy for pancreatic cancer. Cancer letters, 407, 57–65.

Gieniec, K. A., Butler, L. M., Worthley, D. L., & Woods, S. L. (2019). Cancer-associated fibroblasts—heroes or villains?. British journal of cancer, 121(4), 293–302.

Gorvel, L., & Olive, D. (2020). Targeting the “PVR–TIGIT axis” with immune checkpoint therapies. F1000Research, 9.

Griffith, M., Spies, N. C., Krysiak, K., McMichael, J. F., Coffman, A. C., Danos, A. M., … & Barnell, E. K. (2017). CIViC is a community knowledgebase for expert crowdsourcing the clinical interpretation of variants in cancer. Nature genetics, 49(2), 170–174.

Hafemeister, C., & Satija, R. (2019). Normalization and variance stabilization of single-cell RNA-seq data using regularized negative binomial regression. Genome biology, 20(1), 1–15.

Hargadon, K. M. (2016). Dysregulation of TGF β1 activity in cancer and its influence on the quality of anti-tumor immunity. Journal of clinical medicine, 5(9), 76.

Helms, E., Onate, M. K., & Sherman, M. H. (2020). Fibroblast heterogeneity in the pancreatic tumor microenvironment. Cancer Discovery, 10(5), 648–656.

Hosein, A. N., Brekken, R. A., & Maitra, A. (2020). Pancreatic cancer stroma: an update on therapeutic targeting strategies. Nature Reviews Gastroenterology & Hepatology, 1-19.

Huang, Y., Lin, D., & Taniguchi, C. M. (2017). Hypoxia inducible factor (HIF) in the tumor microenvironment: friend or foe?. Science China Life Sciences, 60(10), 1114–1124.

Ilic, M., & Ilic, I. (2016). Epidemiology of pancreatic cancer. World journal of gastroenterology, 22(44), 9694.

Kamburov, A., Stelzl, U., Lehrach, H., & Herwig, R. (2013). The ConsensusPathDB interaction database: 2013 update. Nucleic acids research, 41(D1), D793–D800.

Kang, J., Hwang, I., Yoo, C., Kim, K. P., Jeong, J. H., Chang, H. M., … & Lee, S. K. (2018). Nab-paclitaxel plus gemcitabine versus FOLFIRINOX as the first-line chemotherapy for patients with metastatic pancreatic cancer: retrospective analysis. Investigational New Drugs, 36(4), 732–741.

Karczewski, K., & Francioli, L. (2017). The genome aggregation database (gnomAD). MacArthur Lab.

Klose, R., Krzywinska, E., Castells, M., Gotthardt, D., Putz, E. M., Kantari-Mimoun, C., … & Fandrey, J. (2016). Targeting VEGF-A in myeloid cells enhances natural killer cell responses to chemotherapy and ameliorates cachexia. Nature communications, 7(1), 1–14.

Kopp, J. L., von Figura, G., Mayes, E., Liu, F. F., Dubois, C. L., Morris IV, J. P., … & Hebrok, M. (2012). Identification of Sox9-dependent acinar-to-ductal reprogramming as the principal mechanism for initiation of pancreatic ductal adenocarcinoma. Cancer cell, 22(6), 737–750.

Koukourakis, M. I., Giatromanolaki, A., Sivridis, E., Simopoulos, C., Turley, H., Talks, K., … & Tumour and Angiogenesis Research Group. (2002). Hypoxia-inducible factor (HIF1A and HIF2A), angiogenesis, and chemoradiotherapy outcome of squamous cell head-and-neck cancer. International Journal of Radiation Oncology* Biology* Physics, 53(5), 1192–1202.

Kraman, M., Bambrough, P. J., Arnold, J. N., Roberts, E. W., Magiera, L., Jones, J. O., … & Fearon, D. T. (2010). Suppression of antitumor immunity by stromal cells expressing fibroblast activation protein–α. Science, 330(6005), 827–830.

Lasher, D., Szabó, A., Masamune, A., Chen, J. M., Xiao, X., Whitcomb, D. C., … & Issarapu, P. (2019). Protease-sensitive pancreatic lipase (PNLIP) variants are associated with early onset chronic pancreatitis. The American journal of gastroenterology, 114(6), 974.

Le Maréchal, C., Masson, E., Chen, J. M., Morel, F., Ruszniewski, P., Levy, P., & Férec, C. (2006). Hereditary pancreatitis caused by triplication of the trypsinogen locus. Nature genetics, 38(12), 1372–1374.

Li, Q., Wang, H., Zogopoulos, G., Shao, Q., Dong, K., Lv, F., … & Gao, Z. H. (2016). Reg proteins promote acinar-to-ductal metaplasia and act as novel diagnostic and prognostic markers in pancreatic ductal adenocarcinoma. Oncotarget, 7(47), 77838.

Liekens, S., Schols, D., & Hatse, S. (2010). CXCL12-CXCR4 axis in angiogenesis, metastasis and stem cell mobilization. Current pharmaceutical design, 16(35), 3903–3920.

Looi, C. K., Chung, F. F. L., Leong, C. O., Wong, S. F., Rosli, R., & Mai, C. W. (2019). Therapeutic challenges and current immunomodulatory strategies in targeting the immunosuppressive pancreatic tumor microenvironment. Journal of Experimental & Clinical Cancer Research, 38(1), 162.

Makohon-Moore, A. P., Matsukuma, K., Zhang, M., Reiter, J. G., Gerold, J. M., Jiao, Y., … & Hruban, R. H. (2018). Precancerous neoplastic cells can move through the pancreatic ductal system. Nature, 561(7722), 201–205.

Maurer, C., Holmstrom, S. R., He, J., Laise, P., Su, T., Ahmed, A., … & Genkinger, J. M. (2019). Experimental microdissection enables functional harmonisation of pancreatic cancer subtypes. Gut, 68(6), 1034–1043.

McGuigan, A., Kelly, P., Turkington, R. C., Jones, C., Coleman, H. G., & McCain, R. S. (2018). Pancreatic cancer: A review of clinical diagnosis, epidemiology, treatment and outcomes. World journal of gastroenterology, 24(43), 4846.

Mertins, P., Tang, L. C., Krug, K., Clark, D. J., Gritsenko, M. A., Chen, L., … & Petyuk, V. A. (2018). Reproducible workflow for multiplexed deep-scale proteome and phosphoproteome analysis of tumor tissues by liquid chromatography–mass spectrometry. Nature protocols, 13(7), 1632–1661.

Moffitt, R. A., Marayati, R., Flate, E. L., Volmar, K. E., Loeza, S. G. H., Hoadley, K. A., … & Smyla, J. K. (2015). Virtual microdissection identifies distinct tumor-and stroma-specific subtypes of pancreatic ductal adenocarcinoma. Nature genetics, 47(10), 1168.

Morrison, A. H., Byrne, K. T., & Vonderheide, R. H. (2018). Immunotherapy and prevention of pancreatic cancer. Trends in cancer, 4(6), 418–428.

Murai, J., Thomas, A., Miettinen, M., & Pommier, Y. (2019). Schlafen 11 (SLFN11), a restriction factor for replicative stress induced by DNA-targeting anti-cancer therapies. Pharmacology & Therapeutics, 201, 94–102.

Murphy, S. J., Hart, S. N., Lima, J. F., Kipp, B. R., Klebig, M., Winters, J. L., … & Scherer, S. E. (2013). Genetic alterations associated with progression from pancreatic intraepithelial neoplasia to invasive pancreatic tumor. Gastroenterology, 145(5), 1098–1109.

Nieuwenhuis, T. O., Yang, S. Y., Verma, R. X., Pillalamarri, V., Arking, D. E., Rosenberg, A. Z., … & Halushka, M. K. (2020). Consistent RNA sequencing contamination in GTEx and other data sets. Nature communications, 11(1), 1–10.

Niknami, Z., Eslamifar, A., Emamirazavi, A., Ebrahimi, A., & Shirkoohi, R. (2017). The association of vimentin and fibronectin gene expression with epithelial-mesenchymal transition and tumor malignancy in colorectal carcinoma. EXCLI journal, 16, 1009.

Peitzsch, C., Cojoc, M., Kurth, I., & Dubrovska, A. (2015). Implications of CXCR4/CXCL12 Interaction for Cancer Stem Cell Maintenance and Cancer Progression. In Cancer Stem Cells: Emerging Concepts and Future Perspectives in Translational Oncology (pp. 89–130). Springer, Cham.

Peng, J., Sun, B. F., Chen, C. Y., Zhou, J. Y., Chen, Y. S., Chen, H., … & Yang, Y. (2019). Single-cell RNA-seq highlights intra-tumoral heterogeneity and malignant progression in pancreatic ductal adenocarcinoma. Cell research, 29(9), 725–738.

Pereira, B. A., Vennin, C., Papanicolaou, M., Chambers, C. R., Herrmann, D., Morton, J. P., … & Timpson, P. (2019). CAF subpopulations: a new reservoir of stromal targets in pancreatic Cancer. Trends in cancer, 5(11), 724–741.

Plava, J., Cihova, M., Burikova, M., Matuskova, M., Kucerova, L., & Miklikova, S. (2019). Recent advances in understanding tumor stroma-mediated chemoresistance in breast cancer. Molecular cancer, 18(1), 67.

Pu, N., Lou, W., & Yu, J. (2019). PD-1 immunotherapy in pancreatic cancer: current status. Journal of Pancreatology, 2(1), 6–10.

Qadir, M. M. F., Álvarez-Cubela, S., Klein, D., van Dijk, J., Muñiz-Anquela, R., Moreno-Hernández, Y. B., … & Díaz, Á. (2020). Single-cell resolution analysis of the human pancreatic ductal progenitor cell niche. Proceedings of the National Academy of Sciences, 117(20), 10876–10887.

Raphael, B. J., Hruban, R. H., Aguirre, A. J., Moffitt, R. A., Yeh, J. J., Stewart, C., … & Gabriel, S. B. (2017). Integrated genomic characterization of pancreatic ductal adenocarcinoma. Cancer cell, 32(2), 185–203.

Raphael, K. L., & Willingham, F. F. (2016). Hereditary pancreatitis: current perspectives. Clinical and experimental gastroenterology, 9, 197.

Rawla, P., Sunkara, T., & Gaduputi, V. (2019). Epidemiology of pancreatic cancer: global trends, etiology and risk factors. World journal of oncology, 10(1), 10.

Reches, A., Ophir, Y., Stein, N., Kol, I., Isaacson, B., Amikam, Y. C., … & Seliger, B. (2020). Nectin4 is a novel TIGIT ligand which combines checkpoint inhibition and tumor specificity. Journal for ImmunoTherapy of Cancer, 8(1), e000266.

Ren, B., Cui, M., Yang, G., Wang, H., Feng, M., You, L., & Zhao, Y. (2018). Tumor microenvironment participates in metastasis of pancreatic cancer. Molecular cancer, 17(1), 108.

Saad, A. M., Turk, T., Al-Husseini, M. J., & Abdel-Rahman, O. (2018). Trends in pancreatic adenocarcinoma incidence and mortality in the United States in the last four decades; a SEER-based study. BMC cancer, 18(1), 688.

Sahai, E., Astsaturov, I., Cukierman, E., DeNardo, D. G., Egeblad, M., Evans, R. M., … & Hynes, R. O. (2020). A framework for advancing our understanding of cancer-associated fibroblasts. Nature Reviews Cancer, 1–13.

Sanchez-Vega, F., Mina, M., Armenia, J., Chatila, W. K., Luna, A., La, K. C., … & Chakravarty, D. (2018). Oncogenic signaling pathways in the cancer genome atlas. Cell, 173(2), 321–337.

Schnurr, M., Duewell, P., Bauer, C., Rothenfusser, S., Lauber, K., Endres, S., & Kobold, S. (2015). Strategies to relieve immunosuppression in pancreatic cancer. Immunotherapy, 7(4), 363–376.

Scott, A. D., Huang, K. L., Weerasinghe, A., Mashl, R. J., Gao, Q., Martins Rodrigues, F., … & Ding, L. (2019). CharGer: clinical Characterization of Germline variants. Bioinformatics, 35(5), 865–867.

Semenza, G. L. (2003). Targeting HIF-1 for cancer therapy. Nature reviews cancer, 3(10), 721–732.

Siegel, R. L., Miller, K. D., & Jemal, A. (2020). Cancer statistics, 2020. CA: a cancer journal for clinicians, 70(1), 7–30.

Storz, P. (2017). Acinar cell plasticity and development of pancreatic ductal adenocarcinoma. Nature reviews Gastroenterology & hepatology, 14(5), 296–304.

Stotz, M., Barth, D. A., Riedl, J. M., Asamer, E., Klocker, E. V., Kornprat, P., … & Gerger, A. (2020). The Lipase/Amylase Ratio (LAR) in Peripheral Blood Might Represent a Novel Prognostic Marker in Patients with Surgically Resectable Pancreatic Cancer. Cancers, 12(7), 1798.

Tickle, T., TI, G. C., Brown, M., & Haas, B. (2019). inferCNV of the Trinity CTAT Project. Klarman Cell Observatory, Broad Institute of MIT and Harvard, Cambridge, MA, USA.

Torres, C., & Grippo, P. J. (2018). Pancreatic cancer subtypes: a roadmap for precision medicine. Annals of medicine, 50(4), 277–287.

Uzunparmak, B., & Sahin, I. H. (2019). Pancreatic cancer microenvironment: a current dilemma. Clinical and translational medicine, 8(1), 2.

Valkenburg, K. C., de Groot, A. E., & Pienta, K. J. (2018). Targeting the tumour stroma to improve cancer therapy. Nature reviews Clinical oncology, 15(6), 366–381.

Valque, H., Gouyer, V., Gottrand, F., & Desseyn, J. L. (2012). MUC5B leads to aggressive behavior of breast cancer MCF7 cells. PLoS One, 7(10), e46699.

Wang, D. Y., Salem, J. E., Cohen, J. V., Chandra, S., Menzer, C., Ye, F., … & Rathmell, W. K. (2018). Fatal toxic effects associated with immune checkpoint inhibitors: a systematic review and meta-analysis. JAMA oncology, 4(12), 1721–1728.

Wang, F., Wang, Q., Mohanty, V., Liang, S., Dou, J., Han, J., … & Chen, K. (2020). Single-cell copy number lineage tracing enabling gene discovery. bioRxiv.

Whitcomb, D. C., LaRusch, J., Krasinskas, A. M., Klei, L., Smith, J. P., Brand, R. E., … & Guda, N. M. (2012). Common genetic variants in the CLDN2 and PRSS1-PRSS2 loci alter risk for alcohol-related and sporadic pancreatitis. Nature genetics, 44(12), 1349–1354.

Wilkerson, M. D., & Hayes, D. N. (2010). ConsensusClusterPlus: a class discovery tool with confidence assessments and item tracking. Bioinformatics, 26(12), 1572–1573.

Xu, Y., Liu, J., Nipper, M., & Wang, P. (2019). Ductal vs. acinar? Recent insights into identifying cell lineage of pancreatic ductal adenocarcinoma. Annals of pancreatic cancer, 2.

Yoshihara, K., Shahmoradgoli, M., Martínez, E., Vegesna, R., Kim, H., Torres-Garcia, W., … & Carter, S. L. (2013). Inferring tumour purity and stromal and immune cell admixture from expression data. Nature communications, 4(1), 1–11.

Zhang, W., & Xu, J. (2017). DNA methyltransferases and their roles in tumorigenesis. Biomarker research, 5(1), 1–8.

Zhang, Z., Wang, Y., Zhang, J., Zhong, J., & Yang, R. (2018). COL1A1 promotes metastasis in colorectal cancer by regulating the WNT/PCP pathway. Molecular Medicine Reports, 17(4), 5037–5042.

Zimta, A. A., Cenariu, D., Irimie, A., Magdo, L., Nabavi, S. M., Atanasov, A. G., & Berindan-Neagoe, I. (2019). The role of Nrf2 activity in cancer development and progression. Cancers, 11(11), 1755.

