## Supplementary figures and images for "Spatial drivers and pre-cancer populations collaborate with the microenvironment in untreated and chemo-resistant pancreatic cancer"

### Supplemental Figures

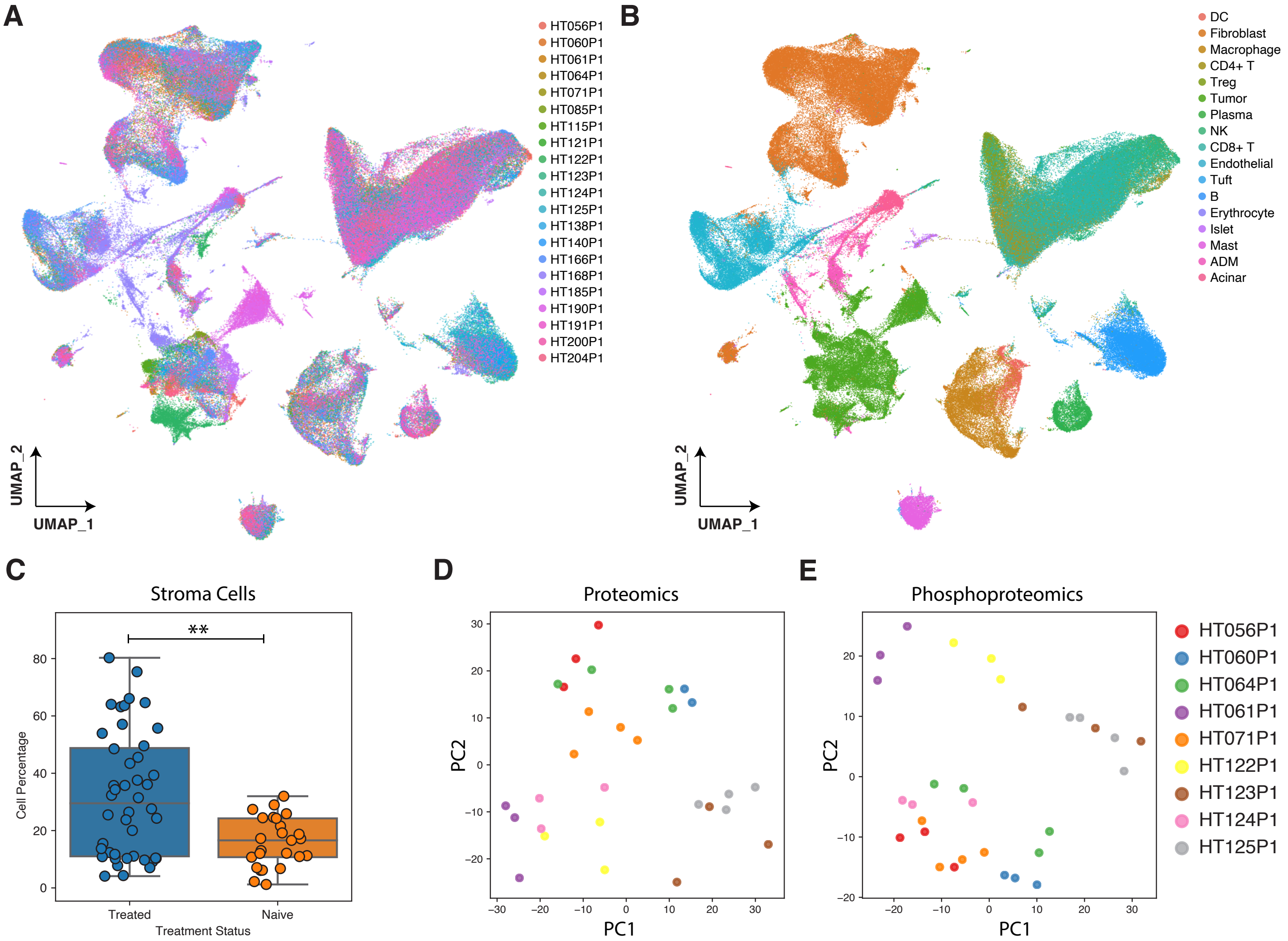

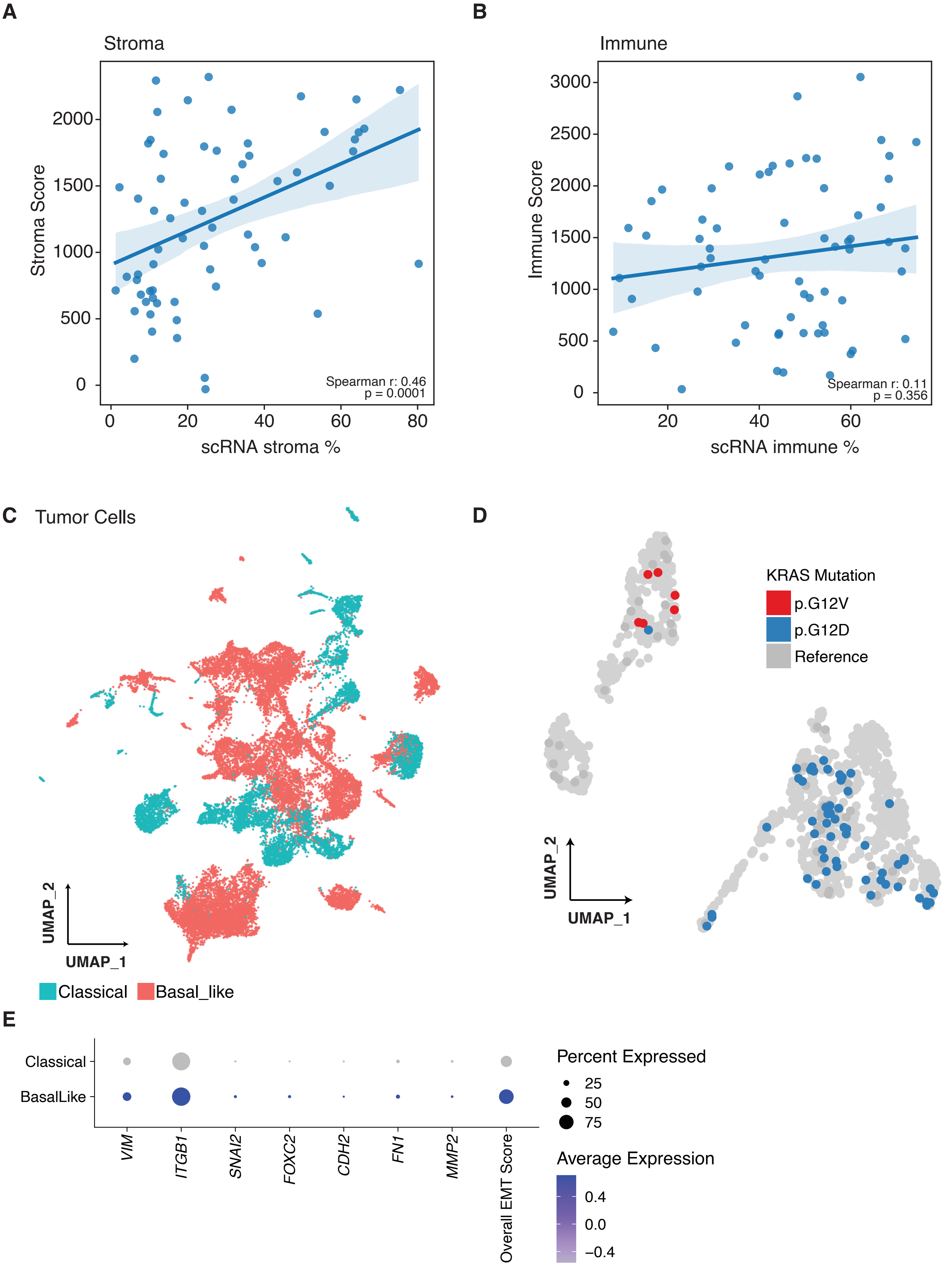

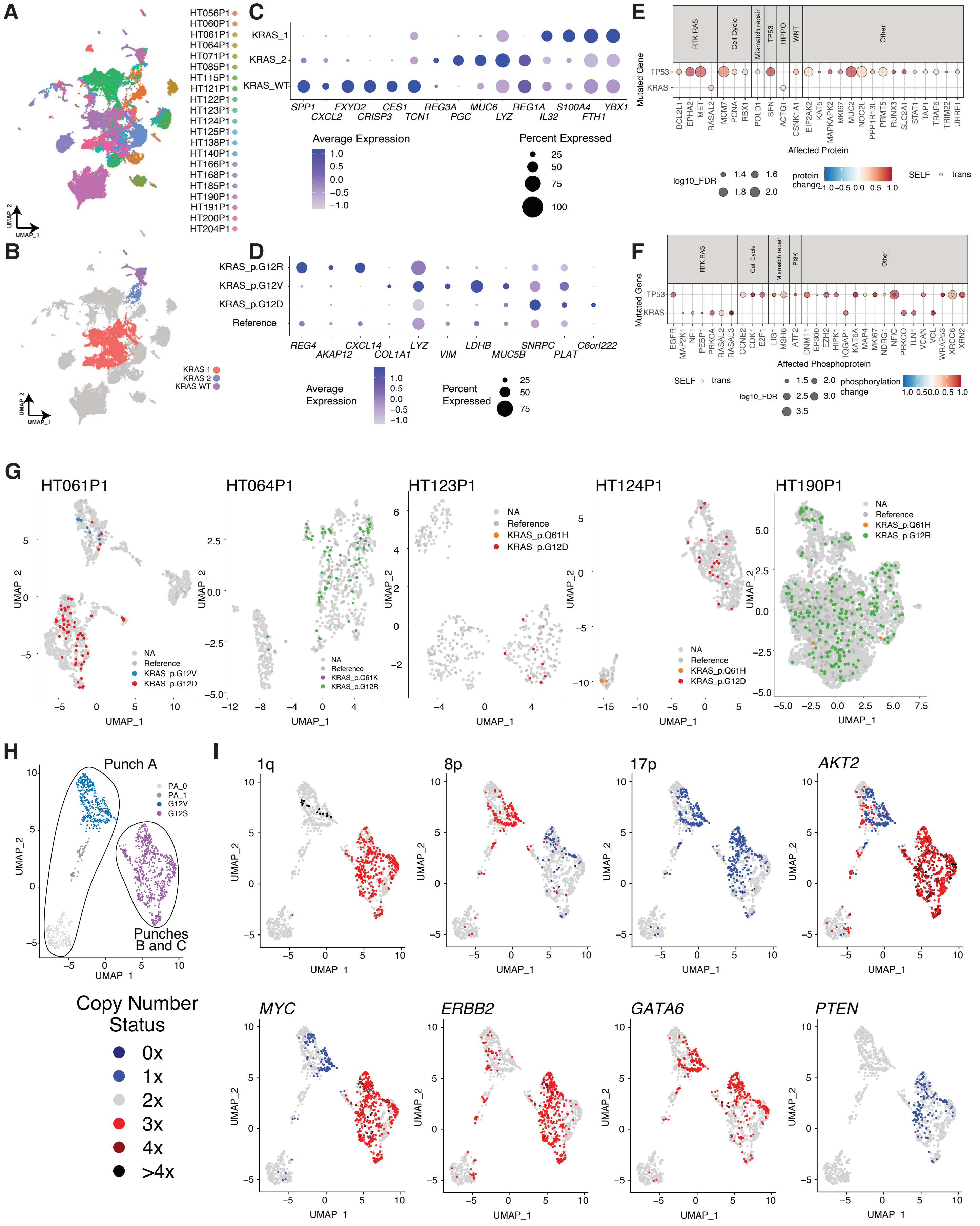

#### Figure S4

A

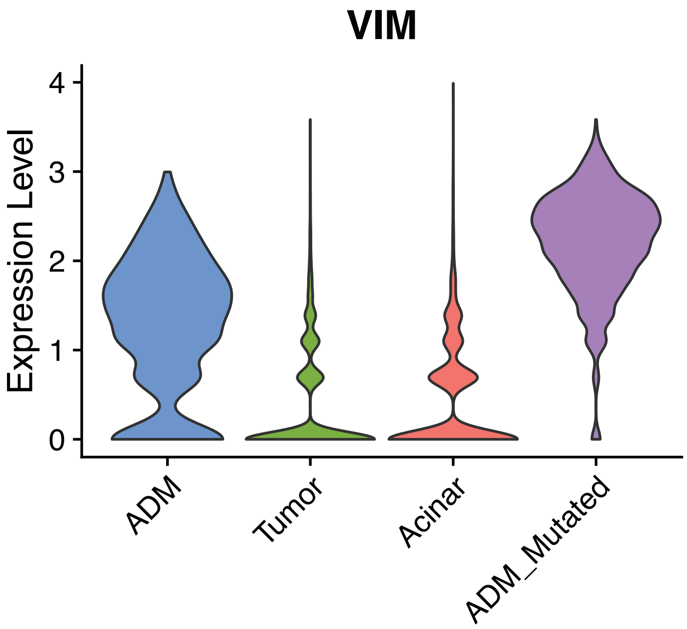


B

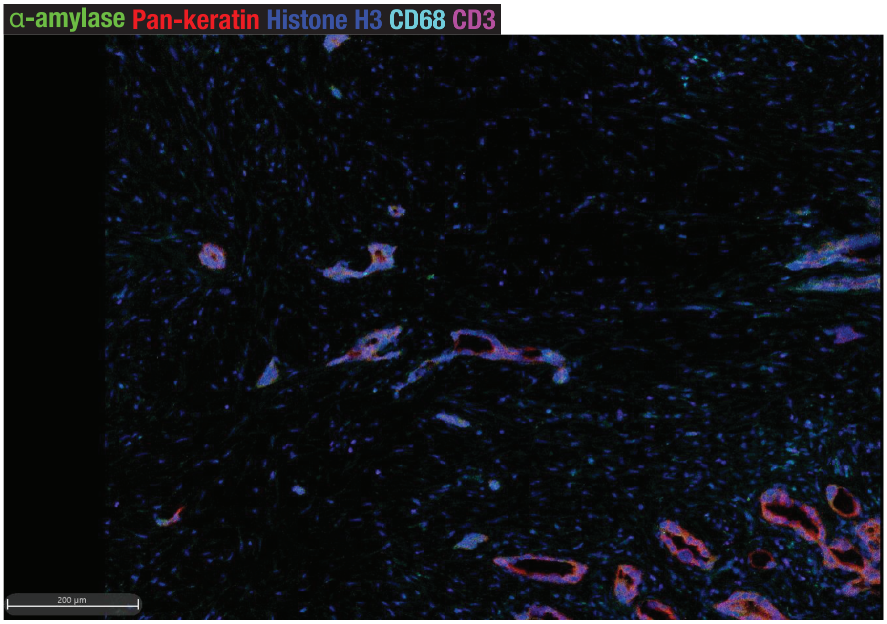


C

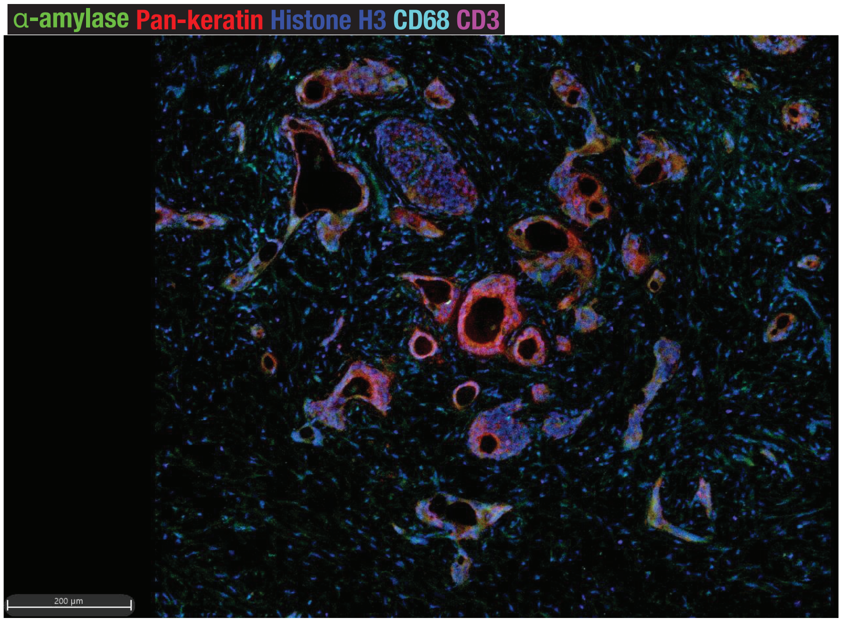

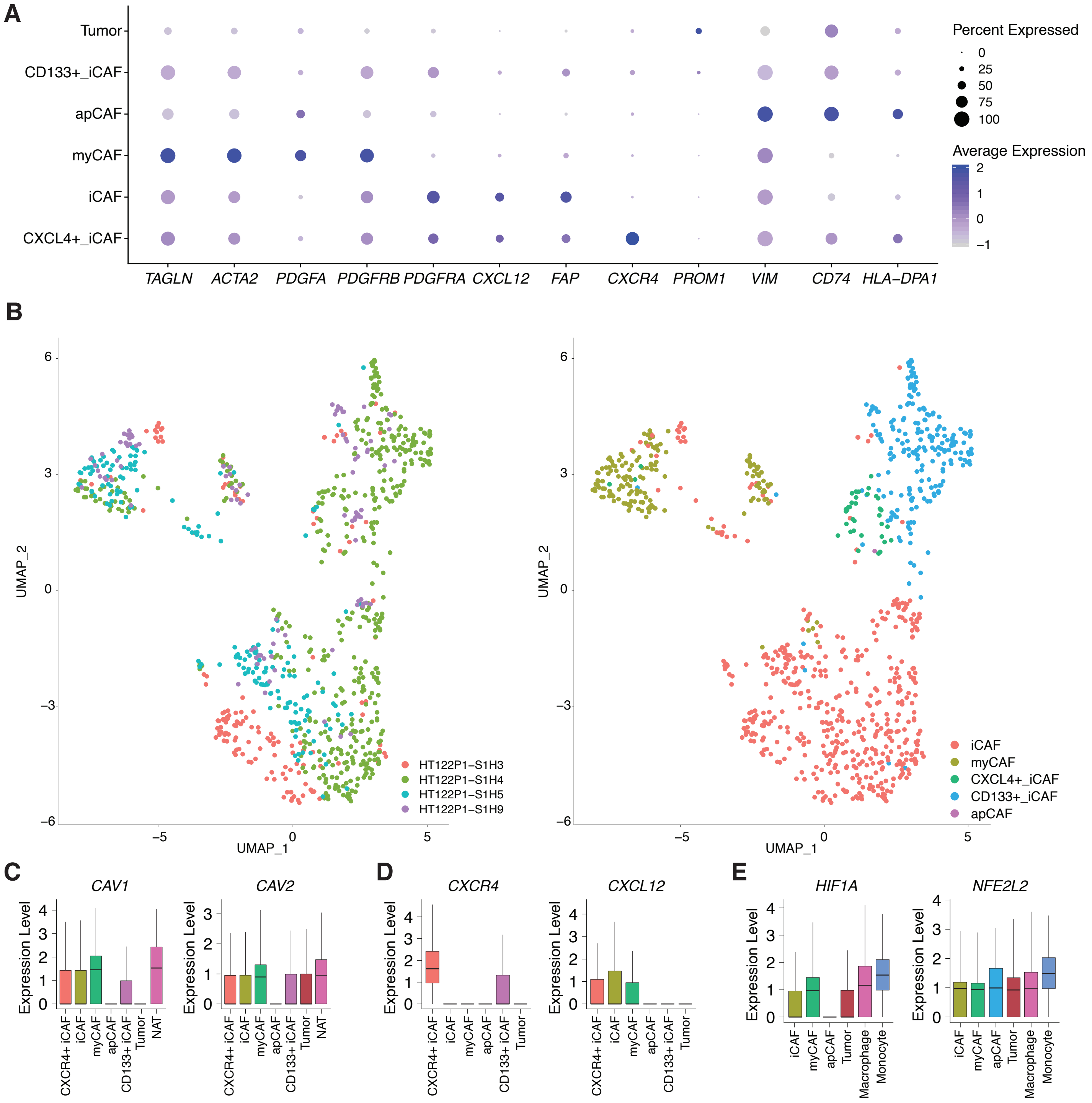

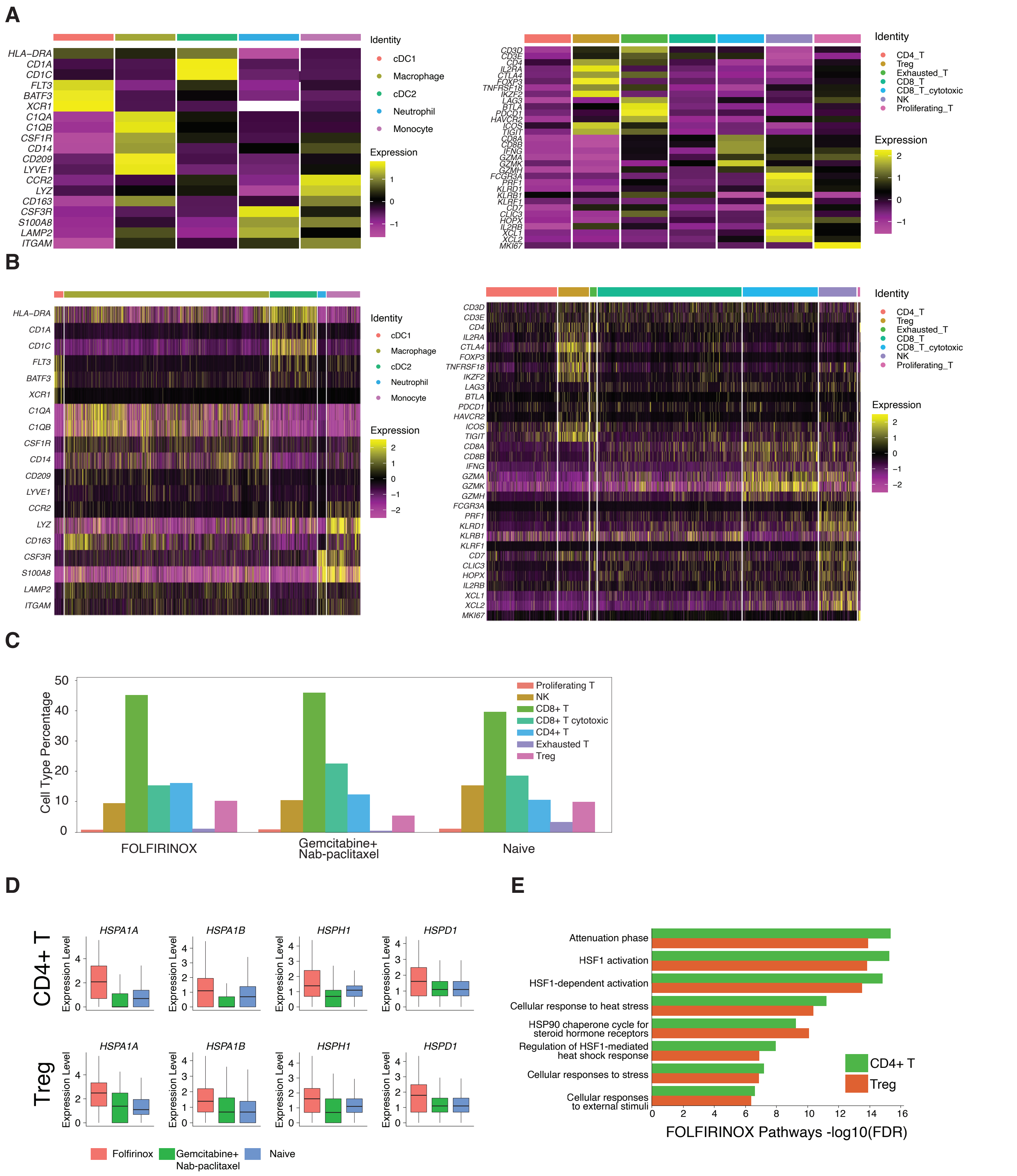

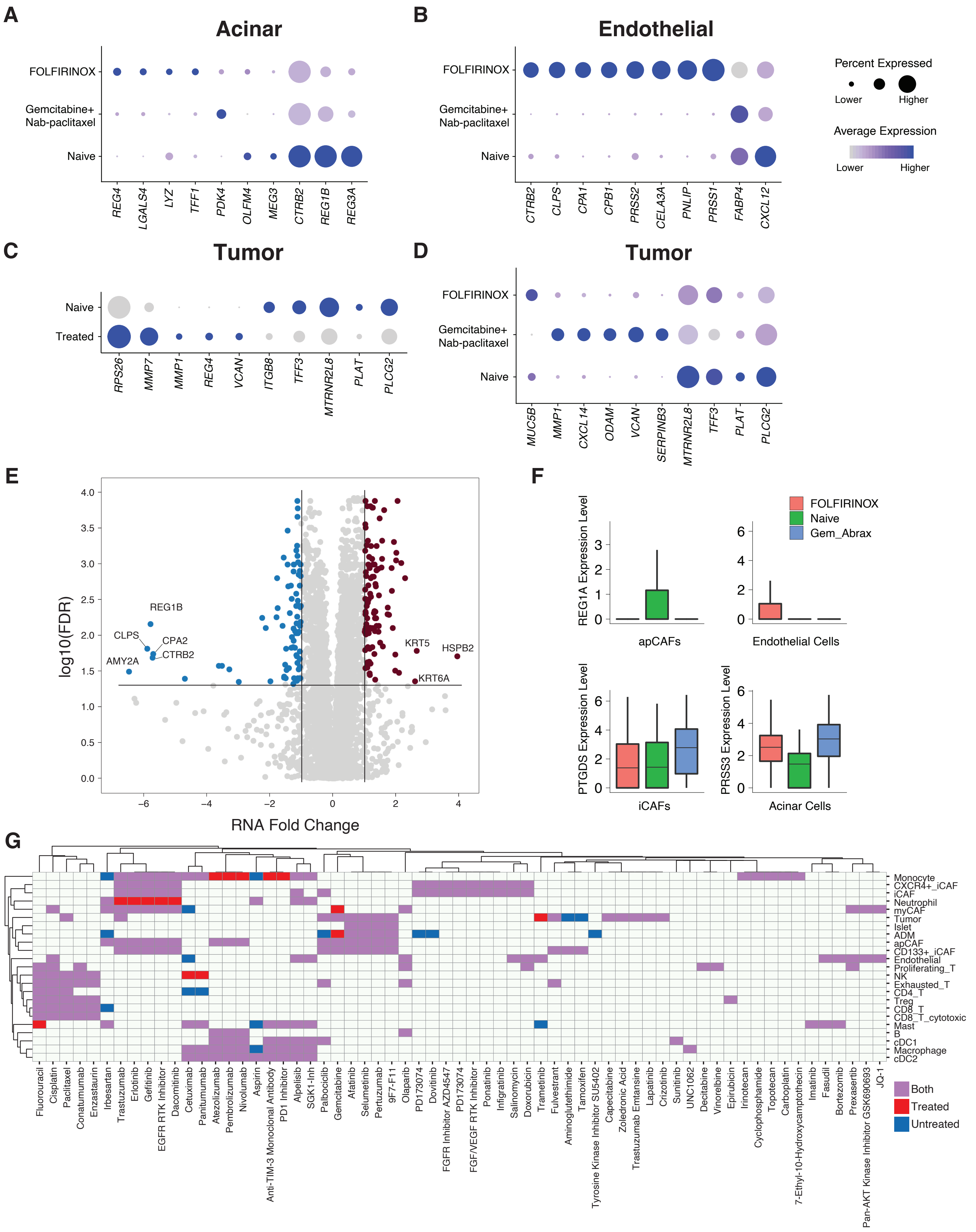
